# Atlas-scale metabolic activities inferred from single-cell and spatial transcriptomics

**DOI:** 10.1101/2025.05.09.653038

**Authors:** Erick Armingol, James Ashcroft, Magda Mareckova, Martin Prete, Valentina Lorenzi, Cecilia Icoresi Mazzeo, Jimmy Tsz Hang Lee, Marie Moullet, Omer Ali Bayraktar, Christian Becker, Krina Zondervan, Luz Garcia-Alonso, Nathan E. Lewis, Roser Vento-Tormo

## Abstract

Metabolism supplies energy, building blocks, and signaling molecules vital for cell function and communication, but methods to directly measure it at single-cell and/or spatial resolutions remain technically challenging and inaccessible for most researchers. Single-cell and spatial transcriptomics offer high-throughput data alternatives with a rich ecosystem of computational tools. Here, we present scCellFie, a computational framework to infer metabolic activities from human and mouse transcriptomic data at single-cell and spatial resolution. Applied to ∼30 million cell profiles, we generated a comprehensive metabolic atlas across human organs, identifying organ– and cell-type-specific activities. In the endometrium, scCellFie reveals metabolic programs contributing to healthy tissue remodeling during the menstrual cycle, with temporal patterns replicated in data from in vitro cultures. We also uncover disease-associated metabolic alterations in endometriosis and endometrial carcinoma, linked to proinflammatory macrophages, and metabolite-mediated epithelial cell communication, respectively. Ultimately, scCellFie provides a scalable toolbox for extracting interpretable metabolic functionalities from transcriptomic data.

## Introduction

Cellular functions are strongly dependent on metabolism, which transforms nutrients into energy and molecules indispensable for cell growth, maintenance of cellular components, and adaptation to challenges such as oxidative stress^1–3^. Through diverse metabolic pathways, cells allocate resources to synthesize macromolecules, including nucleic acids, proteins, and lipids, that also support specialized functions shaping cellular identity^4^. For instance, metabolism is closely integrated with cellular regulation^5^, where metabolites can act as signaling molecules^6^ to influence cell fate decisions^7^, directly linking cellular state to phenotype^8,9^. Disruptions in metabolic processes can contribute to disease^10,11^, as exemplified by cancer, where altered metabolism leads to malignant transformation and supports tumor initiation, growth, and maintenance^12^.

Effective metabolic coordination is fundamental to the function of the female reproductive system, particularly in the uterine lining (endometrium), which is the site of embryo implantation and nutrient provision for early embryonic development. The endometrium undergoes monthly cyclical regeneration in response to ovarian hormones^13^, a process that depends on sustained cellular proliferation and differentiation, hormone biosynthesis, and cellular signaling, all of which rely on proper metabolic function^14^. Disruptions in metabolic processes, often driven by risk factors such as obesity and insulin resistance, are associated with various endometrial disorders, including endometriosis, endometrial carcinoma, infertility, and pregnancy complications^14,15^.

Endometriosis, a chronic inflammatory disease affecting ∼10% of women in reproductive age^16^, is characterised by the growth of endometrial-like tissue outside the uterus (ectopic endometrium). It often leads to debilitating pain and infertility, the latter of which may also arise from abnormalities in the eutopic endometrium. Metabolically, ectopic endometriotic lesions exhibit increased glycolysis resembling the Warburg effect, which supports the aberrant proliferation of ectopic tissue^17^. Similarly, in endometrial carcinoma, the seventh most common cancer in women globally^18^, excess estrogen produced by adipose tissue contributes to hormonal imbalances that drive metabolic reprogramming and promote uncontrolled cellular proliferation^19,20^. However, a comprehensive characterization of which metabolic changes affect specific endometrial cell types and how they contribute to these diseases is still lacking. Advancing our understanding of endometrial metabolism in this context will be key to identify novel therapeutic targets.

Recent advances in single-cell and spatial metabolomics have revolutionized our understanding of cellular metabolism, yet their application to reproductive tissues remains limited^21,22^. This is primarily due to technical challenges that hinder widespread adoption, such as low sensitivity, large sample requirements, metabolite instability, and difficulties in metabolite identification^23–25^. While metabolomics offer the most direct way to quantify small molecules, single-cell RNA sequencing (scRNA-seq) and spatial transcriptomics offers an alternative approach using gene expression as a proxy for inferring metabolic activity. These transcriptomic datasets are more widely available, scalable, comprehensive, and easier to analyze^26,27^, and can complement direct metabolite measurements to gain an scaled-up approximation of the metabolic landscape within cells and tissues.

Bulk and single-cell transcriptomic data have been used to predict metabolic states through several strategies^9,28–35^. Many of these rely on prior knowledge from metabolic databases (e.g. KEGG^36^ and Reactome^37^) and genome-scale metabolic models^38^ (GEMs). Some tools infer pathway-level activity^28–32^, while others mathematically model metabolite conversion rates^33–35^ (metabolic fluxes). Pathway-based methods are easy to interpret, but often depend on gene-set enrichment analysis^30^, which overlooks functional relationships between genes, enzymes, and reactions. The Cellular Functions InferencE (CellFie) method^29^ leverages pathway-based analyses by incorporating these functional relationships to infer the activity of “metabolic tasks”–modules of reactions converting specific metabolites^39^, such as the ATP generation from glucose–but its application remains limited to bulk-resolution data. In contrast, models based on flux balance analysis^40^ (FBA) can accurately estimate reaction-specific fluxes, but often require additional experimental data and some FBA variants require computationally intensive calculations^33,35^, making them impractical for analyzing larger single-cell datasets. Deep learning models scale well to single-cell resolution, yet they compromise interpretability and their accuracy is often validated using paired transcriptomics-metabolomics data points^34^. Thus, there is a pressing need for approaches that combine interpretability, functional relationships of enzymes, and scalability for large single-cell and spatial transcriptomics atlases^41,42^.

Here we present scCellFie, a computational framework that predicts metabolic activities from single-cell and spatial transcriptomics data using the concept of metabolic tasks^39^. By leveraging prior-knowledge from GEMs^43,44^, scCellFie incorporates new analysis modules, integrates seamlessly with the Scanpy^45^ ecosystem, and addresses key challenges of single-cell data analysis (e.g. scalability and data sparsity). This results in an interpretable, scalable, and comprehensive approach to metabolic analysis. We demonstrate scCellFie’s capabilities by analyzing metabolic activities across ∼30 million cells in the CZI CELLxGENE human cell atlas^27^. Additionally, we discover metabolic pathways involved in the menstrual regenerative cycle of the endometrium in health and disease, including endometriosis and endometrial carcinoma. By combining the analysis of the endometrium of patients with endometriosis and the generation of new spatial transcriptomics data from endometriotic lesions, we further unravel metabolic alterations that contribute to a proinflammatory environment in the endometrium and are preserved in the ectopic lesions. In endometrial carcinoma, we investigate the spatial arrangement of metabolic tasks in the tumor microenvironment, identifying activities correlated with tumorigenesis and linked with malignant cells and their metabolite-mediated intercellular communication.

## Results

### scCellFie infers the activities of metabolic tasks

scCellFie uses prior knowledge of reactions, enzymes, and their encoding genes, obtained from GEMs^43,44^ (Figure 1a), to infer the activity of “metabolic tasks” from single-cell and spatial transcriptomics data. These “tasks” are defined as modules of reactions responsible for the conversion of specific substrate metabolites into target products (see ref.^39^). Unlike existing tools limited to bulk transcriptomics^29^ or small single-cell datasets^33^, scCellFie enables efficient and scalable analysis of large cell atlases through integration with the Scanpy ecosystem, facilitating data preprocessing, downstream analysis, and visualization.

**Figure 1.**
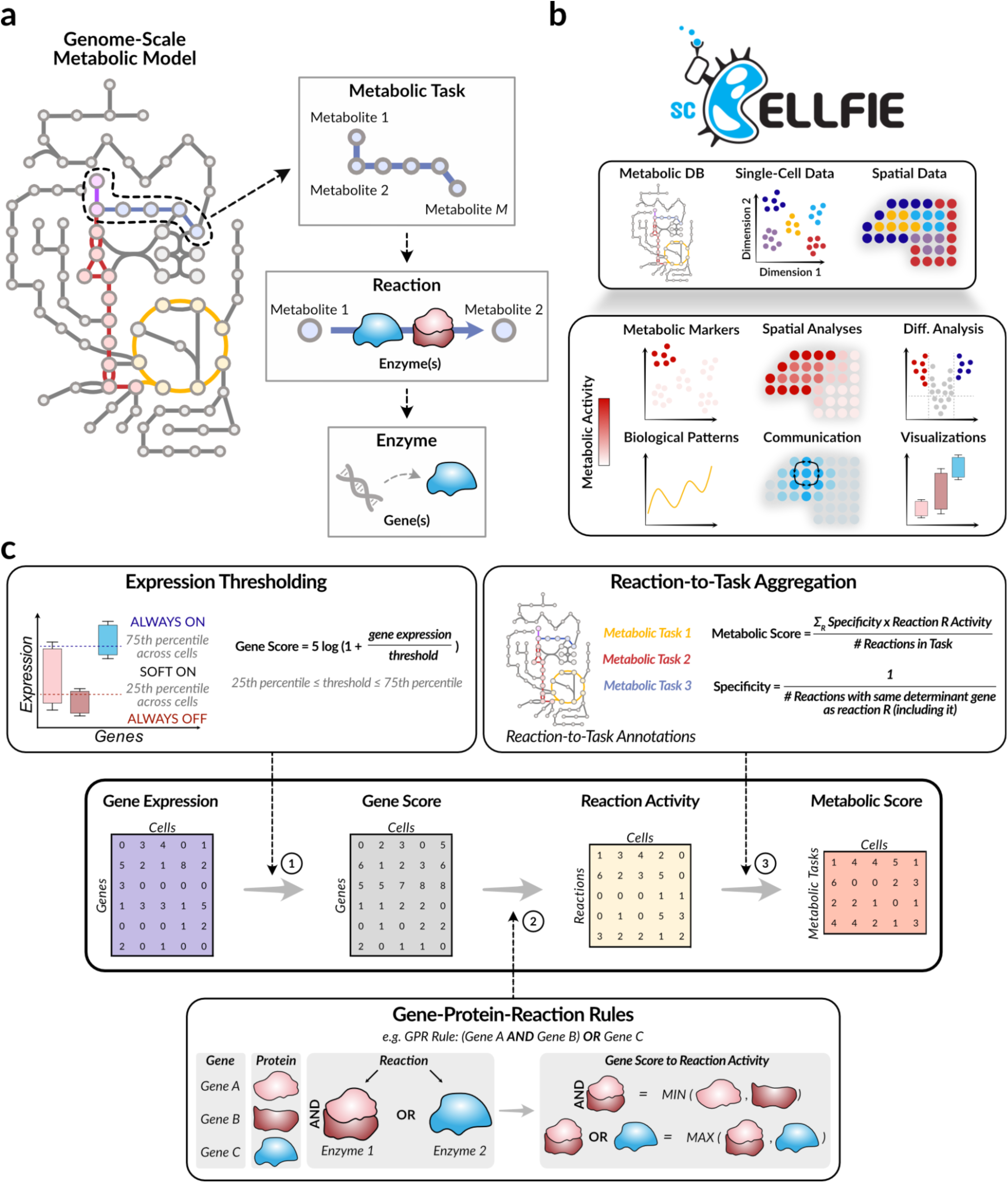
scCellFie is a comprehensive framework to infer, compare, and visualize metabolic task activities from the expression of enzyme-encoding genes. (**a**) Genome-scale metabolic models (GEMs) provide prior knowledge to define metabolic tasks as sets of reactions achieving specific metabolic goals (e.g. transforming one metabolite into another; here, from metabolite 1 to metabolite *M*). Each reaction within a task is catalyzed by isoenzymes, each encoded by one or more genes, with genes corresponding to protein subunits. (**b**) scCellFie uses a metabolic database (Metabolic DB), including expression thresholds and metabolic tasks, to predict metabolic activity from single-cell and spatial data. Users can load scCellFie’s or input their own Metabolic DB. It further enables a series of analyses that make it a comprehensive framework to study metabolism (detailed information in Supplementary Figure 1). (**c**) scCellFie infers metabolic activity by first converting gene expression into gene scores using precomputed or user-defined thresholds that assess enzyme activity potential (step 1). These scores are inputs to compute reaction activities using gene-protein-reaction (GPR) rules, which incorporate functional relationships about enzyme complexes and isoenzymes (step 2). To model metabolic dependencies (bottlenecks) and reaction capacity, it considers the minimum gene score across subunits of a protein complex and the maximum gene score across isoenzymes, respectively. Reaction activities are then aggregated into metabolic task scores (step 3), with specificity weights adjusting for shared reactions among tasks. As colored in purple in panel (**a**), some reactions can be shared by multiple metabolic tasks (in this case by the red and blue tasks). Scores at each of these three main steps can be tracked and used for a range of analyses, including those described in panel (**b**).

scCellFie introduces significant advances in studying metabolic activities at different resolutions (Figure 1b). Building on the strategy for inferring the activity of metabolic tasks, previously introduced in the CellFie method^29^, our tool also incorporates new features that extend its analytical capabilities (Supplementary Figure 1). scCellFie enables single-cell resolution through improvements on optimizing mathematical operations for analysis speed-up and smoothing gene expression to handle data sparsity. Additionally, it goes beyond solely profiling metabolic activity, including modules that use these activities to identify metabolic markers (i.e. metabolic tasks whose activities are overrepresented in specific cell types), condition-specific changes, cell-cell communication (CCC) mediated by non-peptide small molecules, and spatiotemporal patterns (Figure 1b), while offering visualizations that facilitate the interpretation of these analyses. These features make scCellFie a comprehensive framework for studying metabolism.

scCellFie’s workflow consists of three main steps (Figure 1c). First, gene expression data is converted to gene scores using precomputed thresholds. By contextualizing the expression variability across metabolic genes at a cell atlas scale, we can calculate gene-specific thresholds that inform the potential of a gene to be considered “metabolically active”^46^. The scCellFie’s database includes precomputed thresholds across all cells in the CZI CELLxGENE atlas (Supplementary Figure 2), although these thresholds can be optionally customized to better reflect experimental setups. Second, gene scores are translated into reaction activities using gene-protein-reaction (GPR) rules, which capture the functional relationships of enzymes by linking genes, proteins, and reactions^46^. Unlike pathway-based approaches that evaluate gene-set enrichment, GPR rules enable scCellFie to account for enzymes working as protein complexes (where enzymatic activity depends on the presence of all protein subunits) and isoenzymes (where alternative enzymes catalyze the same reaction). Hence, scCellFie can identify activity bottlenecks and reflect biochemical dependencies in the inferred metabolic activity. Third, reaction activities are aggregated into task-specific metabolic scores, and weighted to account for reaction specificity, distributing the activities of determinant genes (i.e. those whose score dominates the AND/OR operations in a reaction’s GPR rule) proportionally across their corresponding reactions.

In addition to precomputed thresholds, the scCellFie database includes 218 metabolic tasks for humans and 203 for mice (Figure 2a). The latter is important to enable the analysis of metabolic activities in animal models of human diseases and in vivo perturbations. Human and mouse tasks cover seven core functions^39^ and five major functions related to protein secretion^47^ (Figure 2b). Within these previously defined tasks, we manually corrected some of their reaction mappings and removed task redundancies (**Methods**). Furthermore, we added two new tasks–tyrosine to melanin and synthesis of GABA–by reutilizing reactions that are already part of the predefined tasks. Importantly, we added a new group of tasks related to hormone metabolism (Figure 2b). In this group, we defined seven new tasks associated with the biosynthesis of sex hormones including testosterone, progesterone, and estrogens (Supplementary Figures 3a,b), using prior knowledge in the GEMs Human1^48^ and Mouse1^49^.

**Figure 2.**
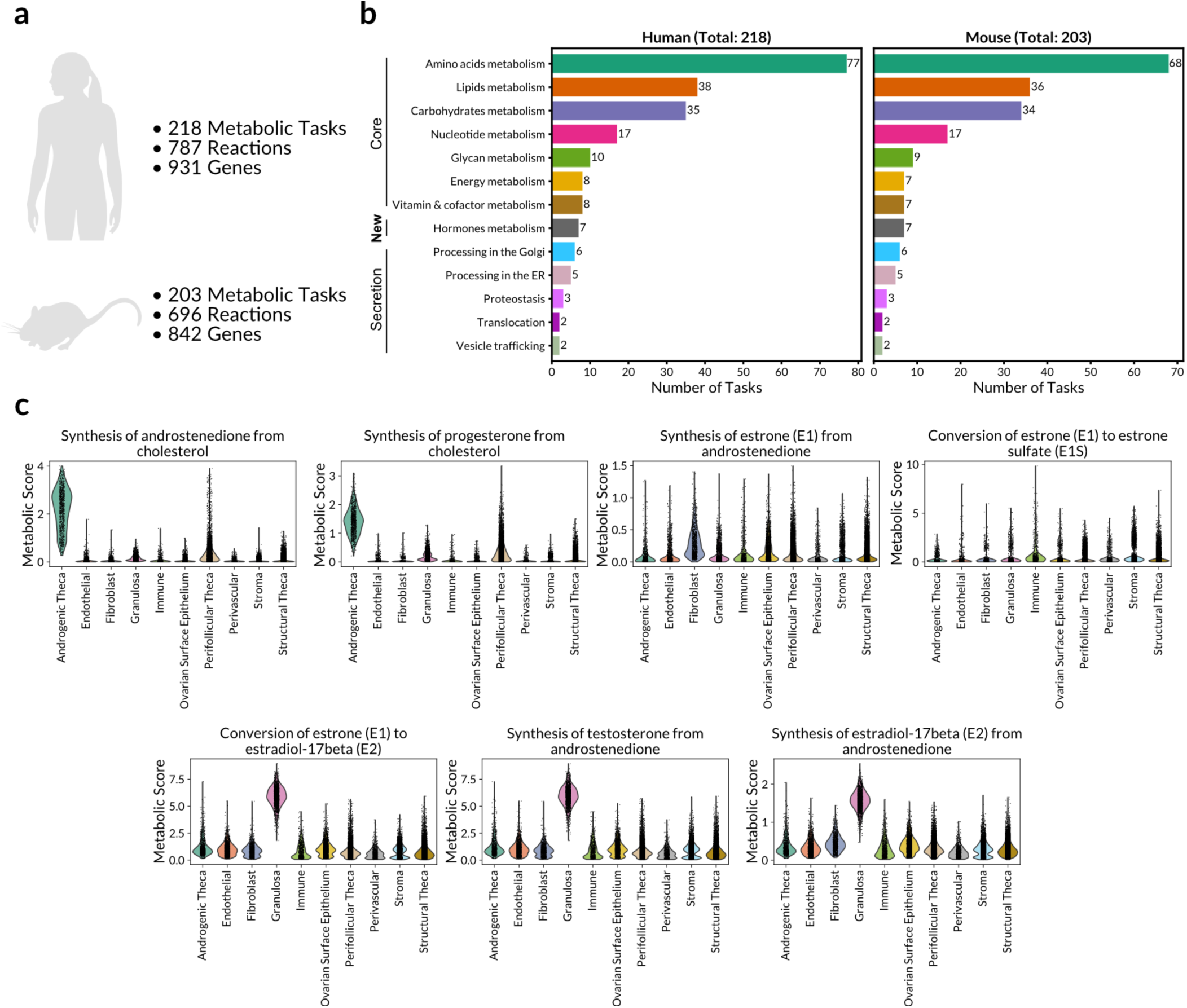
Overview of the metabolic task database in scCellFie and newly added sex-hormone biosynthesis tasks. (**a**) Summary of the number of metabolic tasks, reactions, and genes in the human and mouse databases included in scCellFie. (**b**) Bar plots showing the number of metabolic tasks (x-axis) for each major metabolic function (y-axis) in the human and mouse databases. (**c**) Violin plots summarizing the metabolic activity across single cells within each cell type group for the new sex-hormone biosynthesis tasks included in scCellFie’s database. Metabolic scores representing each task activity were calculated as in Figure 1c from a public dataset of ovaries that includes ∼34k cells covering the diversity of cell types expected to use these tasks^50^ (Supplementary Figure 3). Metabolic scores are not comparable between different tasks, only between cell types for the same task.

To validate the new sex hormone biosynthesis tasks, we evaluated their activity using an ovarian single-cell dataset containing cells involved in complementary biosynthesis steps^50^ (Supplementary Figures 3c,d). Androgenic theca cells are known as the ovarian cells converting cholesterol to progesterone and androstenedione, and scCellFie showed that these cells have the highest activities for these tasks (Figure 2c). Similarly, granulosa cells are known to use androstenedione to synthesize estrone (E1) and estradiol-17β (E2), with a consequent production of testosterone, an intermediate in E2 production (Supplementary Figure 3a); behavior that was captured by scCellFie’s predictions (Figure 2c). scCellFie also inferred that cell types beyond granulosa cells, including fibroblasts and stromal cells, have the potential to produce E1 from androstenedione (Figure 2c), which is consistent with the in vitro production of this hormone by both cell types^51^. Notably, scCellFie revealed an unexpected high activity of immune cells for estrone-to-estrone sulfate conversion (Figure 2c). Given that sulfation is a regulatory mechanism that reduces hormone bioactivity^52,53^, our findings suggest a potential regulation of estrogen homeostasis by the immune cells through hormone sulfation^54^. These results confirm scCellFie’s accuracy in predicting the activity of these new steroid hormone biosynthesis tasks.

### Metabolic activities across cells in the human body

To create a comprehensive metabolic cell atlas covering every cell type across the human body, we ran scCellFie on ∼30 million cells from the CZI CELLxGENE human cell atlas (April 2024 snapshot). Importantly, this CZI atlas includes transcriptomes of cells from 668 published datasets, and harmonizes cell-type labels and metadata, which facilitates cross-organ comparisons. We summarized the metabolic activities and made them available in an online platform (https://www.sccellfie.org) (Figure 3a), where users can explore scCellFie’s predictions and compare metabolic scores between cell types, with their harmonized annotations, across or within organs (Supplementary Figure 4a). Moreover, users can upload to this website their analyses on other datasets for quick visualization and comparison. Ultimately, analyzing the whole human cell atlas demonstrates scCellFie’s scalability and readiness to analyze large datasets.

**Figure 3.**
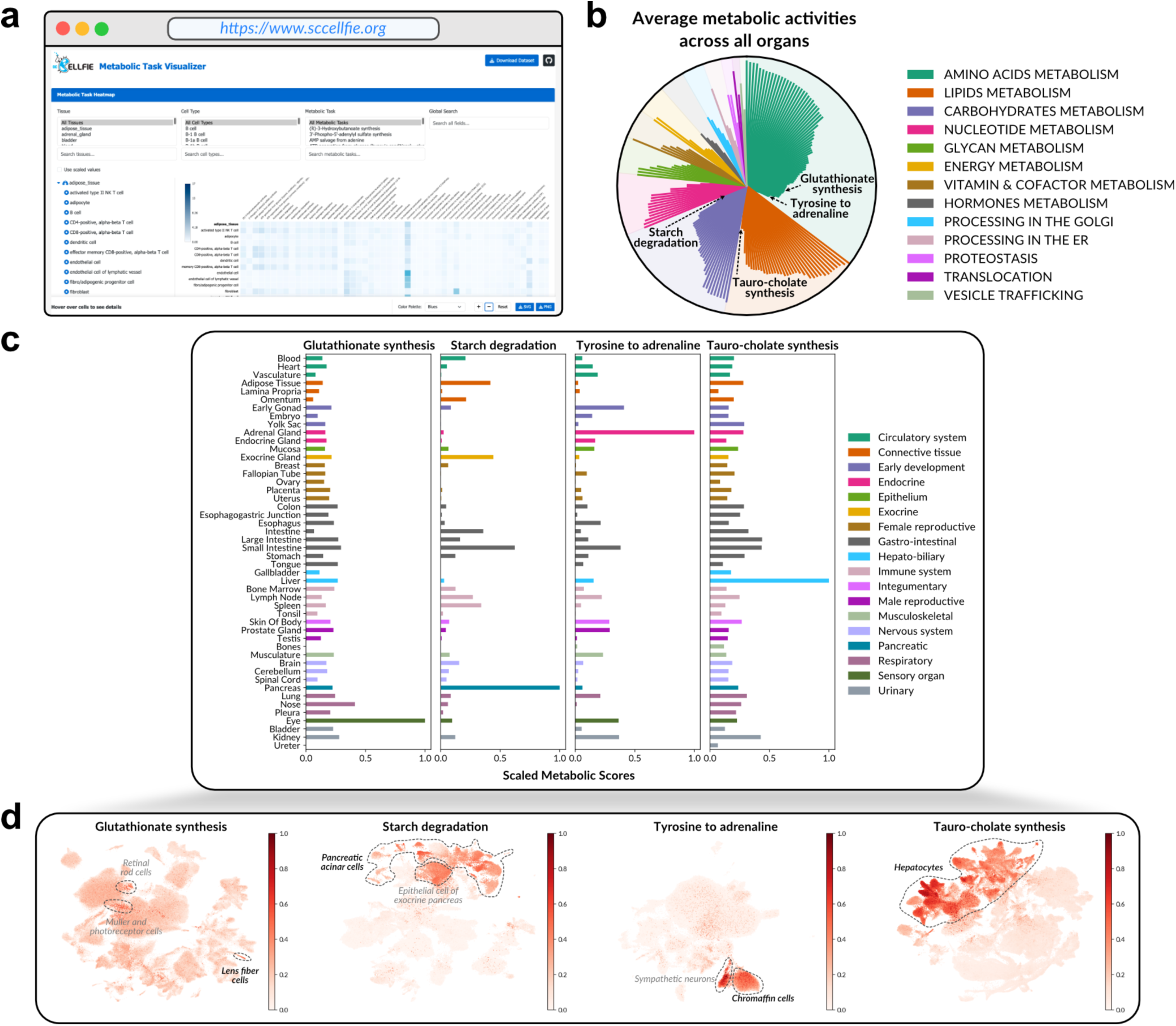
Map of metabolic activities across the CZI CELLxGENE human cell atlas. (**a**) Website to visualize scCellFie’s results. It can either visualize each user’s analysis or summarize the results across every cell type and organ included in the CZI CELLxGENE atlas (April 2024 snapshot). Available online at https://www.sccellfie.org. (**b**) Radial plot showing the average activity of each metabolic task across organs, colored by their major categories. Bars emerging from the center represent the scaled average across organs, where each organ was assigned the maximum metabolic activity found among its cell types. Values range between zero and the highest scaled activity across metabolic tasks (0.68), and tasks are decreasingly sorted. A few metabolic tasks are annotated indicating their corresponding bars in the plot. (**c**) Bar plots showing the maximum activity found across cell types of each organ for the metabolic tasks annotated in (**b**). Cell type activity in each organ was previously aggregated from the single-cell level using the Tukey’s trimean. Organs are colored by their major categories. (**d**) UMAPs for the organs where each metabolic task activity was the highest (see **c**). Cells are colored by their scaled activity for the corresponding metabolic task (inferred activity divided by the maximum value across ∼30M cells). High-activity cells are contoured by a dashed gray line and cell types with known biological functions using these tasks are highlighted. See Supplementary Figure 4 for the same UMAPs with cell type annotations.

We further evaluated scCellFie’s metabolic predictions across organs (Figure 3b). This identified tasks with high overall averages, reflecting they are likely to be central across organs, and others with low averages but high scores in one or few organs. As an example of the latter, glutathionate synthesis was particularly elevated in the eye (Figure 3c), with lens fiber cells among cell types showing the highest activity (Figure 3d), consistent with their known role in producing high amounts of glutathione to maintain lens transparency and counteract oxidative stress^55^. Similarly, starch degradation activity was highest in pancreatic acinar cells (Figure 3d), aligning with their function as the main producers of digestive enzymes such as amylases^56^, whose primary function is to break down starch into nutrients. Adrenaline production was specifically detected in the adrenal gland (Figure 3c), with sympathetic neurons and chromaffin cells showing peak activity (Figure 3d), consistent with these being the main catecholamine-producing cells^57^. The synthesis of taurocholate, a taurine-conjugated bile acid, was predominantly increased in hepatocytes within the liver (Figures 3c,d), accurately reflecting the liver’s role in bile acid production^58^. These examples recapitulating known biology demonstrate scCellFie’s utility for identifying organ– and cell-specific metabolic functions across large-scale atlases such as the CZI CELLxGENE atlas.

### Differences in metabolic activities among endometrial cell types

The human endometrium is a highly dynamic tissue that undergoes considerable changes in its architecture throughout the menstrual cycle (Figures 4a,b), driven by the interplay of the steroid hormones estrogen and progesterone. These changes involve cyclical cellular proliferation and differentiation to support scarless tissue regeneration and key reproductive functions such as embryo implantation, requiring tightly coordinated metabolic adaptations. To understand the endometrium’s metabolic activities and how they support this tissue’s cyclical changes and functions, we applied scCellFie to the Human Endometrial Cell Atlas (HECA) dataset, which provides single-cell transcriptomic profiles of all endometrial populations, including epithelial, mesenchymal, and immune cell lineages, for over 300k cells across different menstrual cycle phases^59,60^.

**Figure 4.**
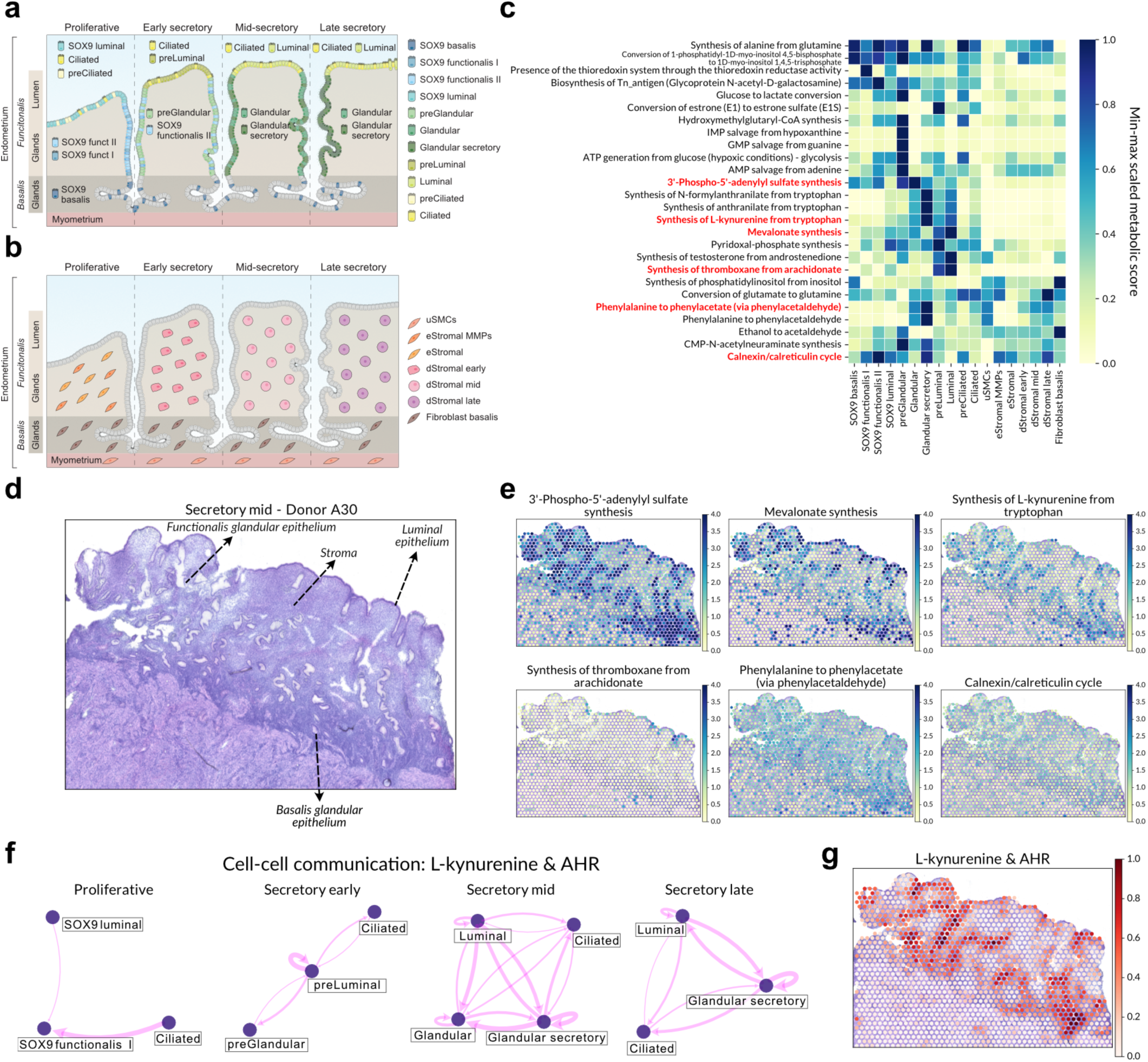
Metabolic markers in the human endometrium at a cell-type and spatial level. (**a**-**b**) Schematic illustrations of the human uterus, highlighting epithelial (**a**) and mesenchymal (**b**) cell compositions during the menstrual cycle. (**c**) Heatmap showing the metabolic tasks identified as markers, after running scCellFie on the HECA dataset, for the cell types displayed in panels (**a**-**b**). Metabolic task activities were aggregated from single cells into cell types using the Tukey’s trimean, and values then min-max scaled based on the scores of each task across the included cell types. In red, we highlight metabolic tasks distinguishing epithelial from mesenchymal cells. (**d**) H&E image of secretory (A30, 152811 slide) endometrium, obtained from a previously published Visium dataset^59^ (**e**) Metabolic markers from (**c**) are shown after running scCellFie on Visium data for the secretory endometrium. (**f**) Cell-cell communication networks, based on the interaction between kynurenine and its receptor AHR, among epithelial and mesenchymal cell types in HECA. Cell types are shown and connected only if their communication scores were above a threshold. Arrow thickness is proportional to their communication score. (**g**) Communication scores inferred for each Visium spot in the secretory endometrium. In this case, scores reflect the co-localization fraction among each spot’s neighborhood based on the metabolic score of the ligand synthesis and the gene expression of the receptor. Radius defining a neighborhood was selected as shown in Supplementary Figure 10a. See Methods for calculation and thresholding of communication scores, either cell-type-or neighborhood-based (panels (**f**) and (**g**), respectively). Panels (**a**) and (**b**) were adapted from Marečková et al (2024)^60^.

scCellFie identified metabolic functions that distinguish specialized roles of epithelial cells (Figure 4c). For instance, epithelial cells showed prominent activity in tasks such as synthesis of 3’-Phospho-5’-adenylyl sulfate (PAPS), which is essential for sulfation reactions regulating hormone bioactivity^61^. Additionally, this synthesis was predicted to be active in regions corresponding to both basalis and functionalis glandular epithelia (Figures 4d,e). scCellFie further predicted a local estrogen inactivation–through PAPS donating a sulfate group to estrone to form estrone sulfate–that was highly specific to the pre-luminal cells in the early secretory phase (Figure 4c). Mevalonate (MVA) synthesis was a task inferred to be active across all epithelial cells, with slightly higher activity in luminal cells (Figures 4c,e). This task supports cholesterol and ubiquinone biosynthesis^62,63^ and regulates cellular proliferation^64^, functions that are essential for maintaining the endometrial epithelia. In addition, synthesis of thromboxane–which promotes platelet aggregation^65^ and modulates uterine contractility^66^–was highly specific to luminal epithelial cells (Figures 4c,e). Thus, this result suggests that luminal cells may rely on thromboxane to regulate hemostasis and tissue shedding, aligning with these cells’ dual roles in implantation and menstruation^67^.

scCellFie predictions can help uncover how different cell types produce and use metabolites for CCC. For example, scCellFie inferred high conversion of tryptophan to L-kynurenine, anthranilate, and N-formylanthranilate in both glandular and luminal epithelial subtypes (Figure 4c). These kynurenine pathway metabolites regulate inflammation and contribute to NAD+ synthesis^68,69^, supporting redox balance and cellular defense against oxidative stress^69^. Notably, scCellFie identified strong communication scores between different epithelial cells across the menstrual cycle (Figure 4f), reflecting a coordinated potential to produce kynurenine and express its receptor, AHR. Spatially, scCellFie predicted increased co-localization of kynurenine synthesis and AHR expression in glands and luminal regions (Figure 4g). These findings suggest that kynurenine-AHR signaling may support endometrial epithelial cells to manage the inflammation and oxidative stress during tissue remodeling^70^.

### Metabolic landscape of the human endometrium across the menstrual cycle

We next used scCellFie to investigate metabolic dynamics across menstrual cycle phases. During the first half of the menstrual cycle (proliferative phase), increasing levels of estrogen, produced in the ovaries, drive tissue repair and proliferation^59^. After ovulation, estrogen levels fluctuate and a rise of progesterone levels, produced by the corpus luteum, induces the secretory phase, promoting differentiation of endometrial fibroblasts into decidualized stroma in preparation to embryo implantation^59^. Hence, based on this knowledge and spatiotemporal dynamics of endometrial cells^59,71^, we first defined the most likely cell-type trajectories across the menstrual cycle (Figure 5a), and then applied scCellFie to identify temporally regulated metabolic tasks. By fitting a generalized additive model (GAM) that is part of the scCelFie toolbox, we identified metabolic tasks changing across the menstrual cycle in glandular epithelial, luminal epithelial, and stromal cells (Figures 5b-d).

**Figure 5.**
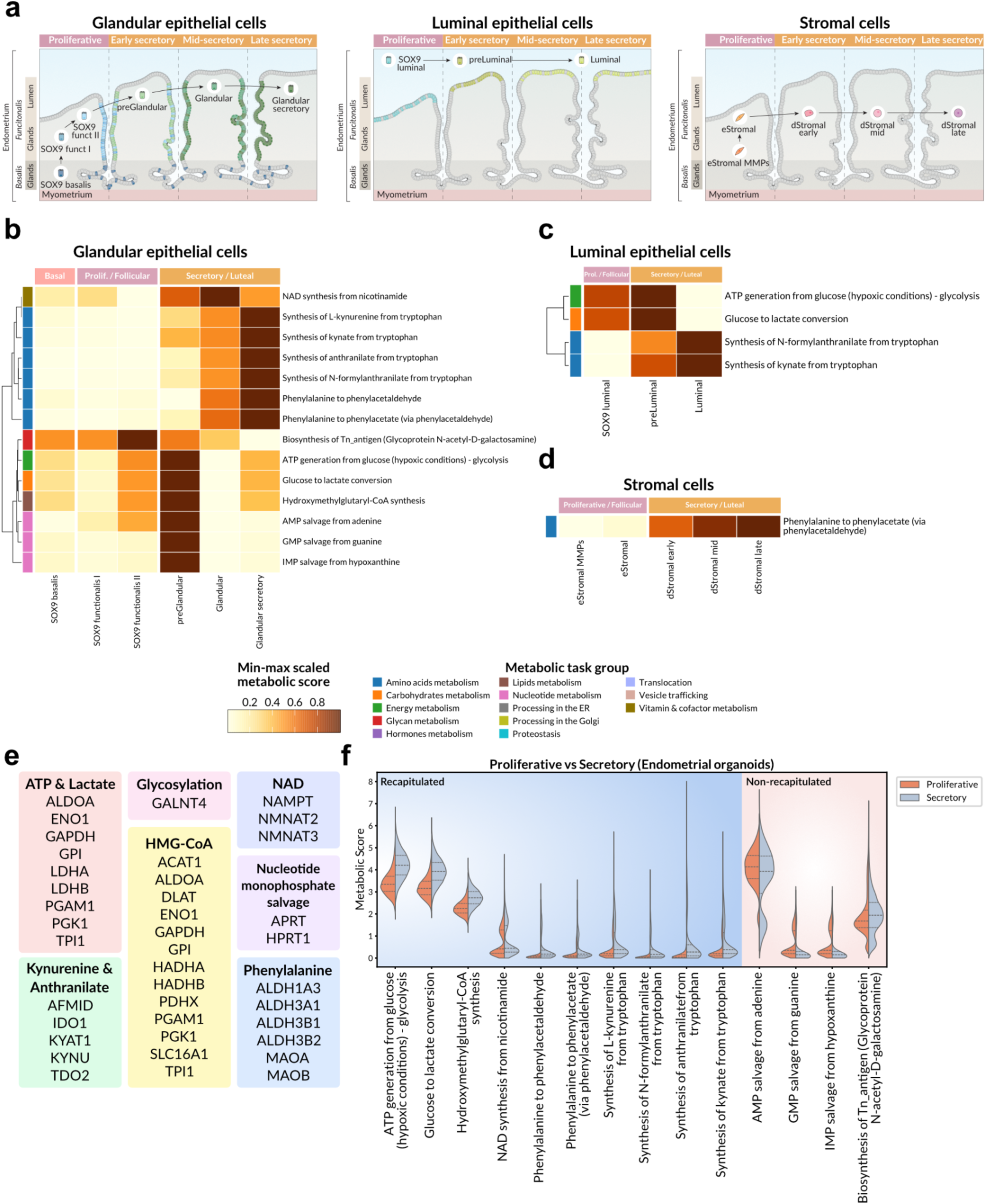
Metabolic changes along cellular trajectories during the menstrual cycle. (**a**) Most likely trajectories of glandular epithelial, luminal epithelial, and stromal cells, based on prior knowledge and ordered according to their progression across the menstrual cycle. (**b**-**d**) Heatmaps summarizing the metabolic tasks changing along the trajectories in (**a**). These dynamic activities were identified using generalized additive models (GAMs) on the activities across single cells. Metabolic scores are shown as the Tukey’s trimeans across single cells within each cell type group. (**e**) Key genes driving the metabolic reactions associated with the tasks in (**d**), grouped by functional categories (bold labels). (**f**) Violin plots comparing the distributions of metabolic activities across epithelial single cells in endometrial organoids, between proliferative and secretory phases of the menstrual cycle. Tasks with similar trends to (**b**-**c**) (peak activity in the same cycle phase) are shown in the blue area, while those following an opposite trend are shown in the red area. Horizontal lines within the violins indicate the first quartile (Q1), median, and third quartile (Q3). All tasks showed significant differences between phases (FDR < 1% after a Wilcoxon rank sum test).

Glandular epithelial cells showed metabolic reprogramming of multiple functions, with 14 tasks showing important temporal variation (Figure 5b). Tasks like the ATP production derived from glycolysis and glucose to lactate conversion peaked during the proliferative-to-secretory transition (Figures 5b,c). Additionally, nucleotide salvage pathways, crucial for energy-efficient nucleotide synthesis^72^, showed similar behavior, with the estrogen-regulated gene APRT^73^ emerging as a key determinant (Figure 5e). The secretory phase, where estrogen levels overall decrease, was further marked by predicted increases in kynurenine production, NAD salvage, and phenylalanine metabolism in glandular epithelial cells. Consistent with this result, kynurenine’s receptor AHR can counterbalance estrogen signaling^74,75^. In the case of luminal cells, only four metabolic tasks were identified by our GAM-based analysis (Figure 5c), suggesting that glandular cells undergo more complex metabolic rewiring. In addition, all of these luminal cells’ tasks are overlapped with glandular epithelial cells (Figure 5b), which may reflect a coordinated metabolic program within the endometrial epithelium.

Decidualized stromal fibroblasts presented high activity in specific metabolic tasks such as the calnexin/calreticulin cycle and phenylalanine conversion to phenylacetate (Figure 4c), with spatial data of early-mid secretory endometrium confirming a wide distribution of these task activities across the tissue (Figures 4d,e). The calnexin/calreticulin cycle^76,77^ active in stromal cells is known to participate in decidualization^78^ and contribute to uterine receptivity^79^. Temporally, the dominant metabolic change in stromal cells as they decidualize during the secretory phase was increased conversion of phenylalanine to phenylacetate (Figure 5d). While this aligns with previous reports of decreased phenylalanine levels during this phase^80^ and known role of progesterone modulating phenylalanine metabolism^81^, our analysis revealed cell-type-specific contributions that go beyond prior knowledge at a tissue level. Additionally, phenylacetate can counteract estrogen activity by inhibiting estrogen response elements (EREs)-associated transcription^82^. Consistent with this, scCellFie predicts an increase in the phenylalanine conversion starting in the early secretory phase (Figure 5d). scCellFie further mapped monoamine oxidases (MAOA and MAOB) as determinant genes of this pathway (Figure 5e). Given that they aid in decidualization and successful embryo implantation^83^, and high phenylalanine levels can impair embryo implantation^84^, this metabolic activity may be central in regulating this amino acid’s abundance after ovulation and preparing the tissue for a potential implantation.

We next used scCellFie to identify metabolic targets for refinement of organoid models. We analyzed publically available single-cell data from epithelial endometrial organoids^59,85^, and evaluated whether organoid response to hormonal stimulation closely matches the metabolic behavior of in vivo epithelial cells. While most cycle-dependent metabolic patterns observed in vivo were successfully reproduced (Figure 5f), confirming the organoids’ utility as metabolic models, we identified specific discrepancies that could inform organoid optimization. Notably, nucleotide metabolism and Tn antigen glycosylation pathways deviated from expected in vivo trends. These tasks’ determinant genes can be easily tracked (Figure 5e), in this case involving GALNT4, APRT, and HPRT1, which highlights potential targets for further improvement of in vitro protocols. Nevertheless, since these tasks peak during the proliferative-to-scretory switch in vivo (Figure 5b), their divergent behavior may also reflect challenges in replicating temporal transitions in organoid models^86^. Overall, scCellFie shows that most in vivo cycle-dependent processes and their temporal transitions are well preserved in endometrial organoids.

### Dysregulation of metabolic tasks in the eutopic and ectopic endometrium of women with endometriosis

Understanding the metabolic changes in endometriosis is key to design new therapeutic strategies that can mitigate these alterations^87^. Hence, we used scCellFie to compare the metabolic activities of eutopic endometrial cells between donors with and without endometriosis. Our tool identified cell-type specific changes among epithelial, mesenchymal and immune cells (Figure 6a), predominantly affecting amino acids, carbohydrate, and lipids metabolism (Figure 6b).

**Figure 6.**
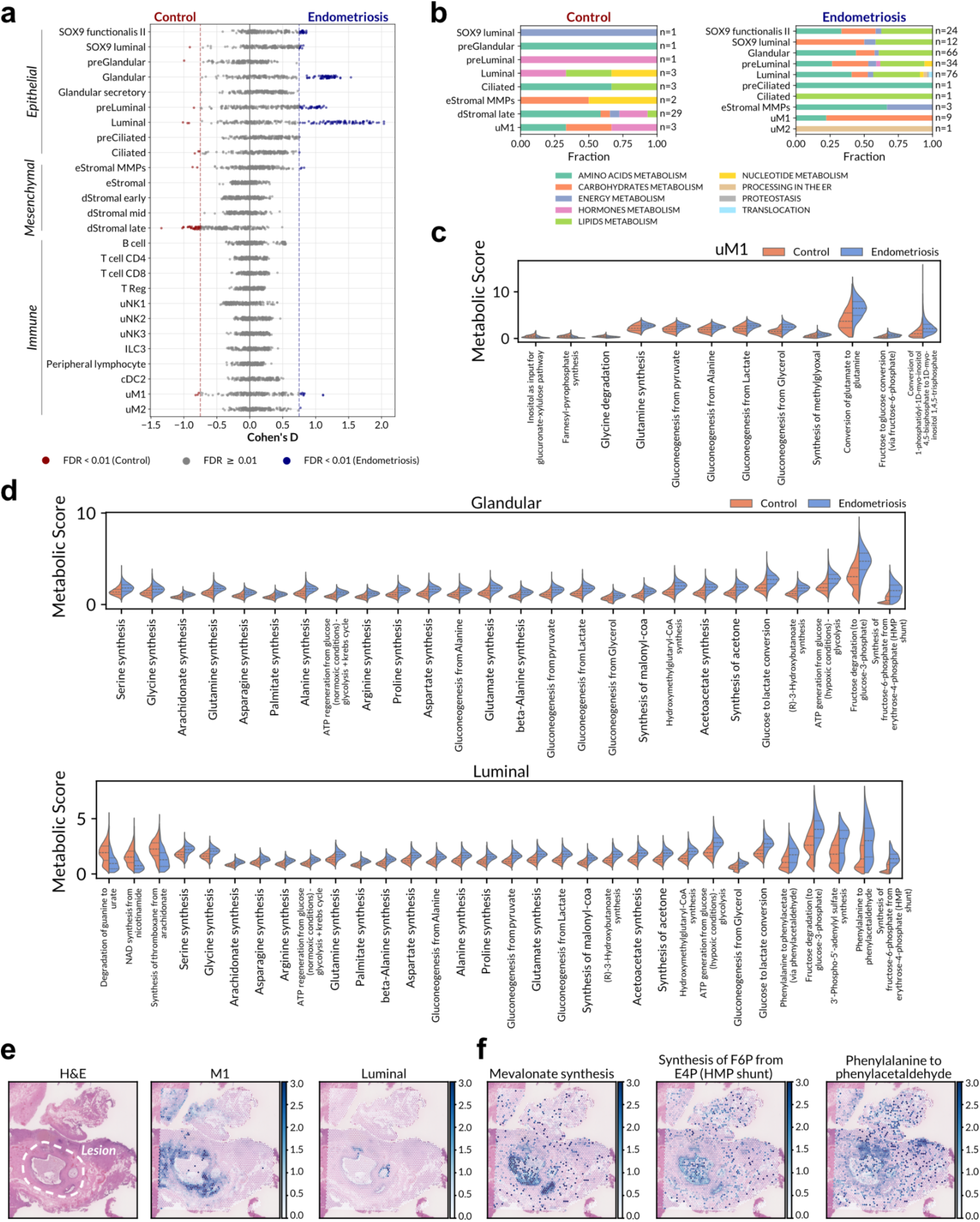
Dysregulated metabolic activities in endometriosis predicted with scCellFie. (**a**) Beeswarm plot of a differential analysis comparing the metabolic scores of each task at a single-cell resolution, per cell type (y-axis), between eutopic endometrium of control and endometriosis donors. The effect size (Cohen’s D, x-axis) is shown for each metabolic task after performing a Wilcoxon rank sum test, and if a task resulted differential between groups (|Cohen’s D| >0.75 and FDR <0.01) it was colored as indicated in the legend. (**b**) Bar plots showing the fraction of differentially altered metabolic tasks per cell type, with higher scores in endometriosis (upper plot) or control (lower plot). (**c**-**d**) Violin plots comparing metabolic score distributions across single cells between control and endometriosis for (**c**) M1 endometrial macrophages, and (**d**) glandular and luminal epithelial cells. In (**d**), a subset of differentially altered tasks is shown (only those with a high average score; a full list of dysregulated metabolic tasks is shown in Supplementary Figure 5). (**e**) Visium data from a deep infiltrating lesion of a donor with endometriosis. H&E image and cell2location scores are displayed, covering the location of M1 macrophages, and luminal epithelial cells. Dashed line encloses the endometriotic lesion in the H&E image. Cell distributions inferred with cell2location were obtained by using single-cell transcriptomes in the HECA dataset as reference. Only cell types with a high score are displayed. (**f**) Metabolic scores are presented for one metabolic marker of epithelial cells (mevalonate synthesis task) and for two differentially altered tasks in luminal epithelial cells (synthesis of F6P from E4P and phenylalanine to phenylacetaldehyde). F6P, fructose-6-phosphate; E4P, erythrose-4-phosphate.

We previously found an imbalance in macrophages in the eutopic endometrium of donors with endometriosis^60^. Consistent with this, scCellFie identified macrophages as the only immune cells with significant metabolic dysregulations in endometriosis (Figure 6a), and specifically revealed that uterine M1 macrophages (uM1) exhibited increased transformation of myo-inositol bisphosphate to myo-inositol triphosphate (Figure 6c), through different isoforms of phospholipases C. This process activates NF-κB signaling^88^, which plays a key role in inflammation, supporting the observation that uM1 from endometriosis patients are more proinflammatory^60^. Additionally, scCellFie inferred an increased production of methylglyoxal in uM1, a reactive metabolite known to induce direct tissue damage and trigger a proinflammatory response^89^.

In addition, we observed a high number of upregulated metabolic activities in glandular and luminal epithelial cells in the mid secretory phase in endometriosis (Figure 6b), including upregulation of malonyl-CoA synthesis, a precursor for fatty acid synthesis and oxidation, along with elevated synthesis of its derived lipids (Supplementary Figure 5). Notably, these cells also showed increased glucose-to-lactate conversion, a central metabolic activity supporting aberrant cellular proliferation^90^. Arachidonate synthesis was also predicted to increase, producing the fatty acid precursor for prostaglandins, key mediators of inflammation. In contrast, stromal cells presented downregulated tasks involved in mitigating inflammation and oxidative stress, including the kynurenine and NAD salvage pathways (Supplementary Figure 5). Collectively, this metabolic shift in epithelial and stromal cells coupled with the metabolic activity in uM1 supports the hypothesis of a proinflammatory environment in the eutopic endometrium of women with endometriosis^60,91^.

We also assessed whether the metabolic activities altered in eutopic endometrium of women with endometriosis are present in ectopic endometrium (also known as endometriotic lesions). We generated spatial transcriptomics data (Visium) of peritoneal endometriotic lesions from two different women (Figure 6e, Supplementary Figure 6), and deconvolved the cell type composition of each Visium spot with cell2location^92^ to identify which endometrial cells are present in the endometriotic lesions. This analysis revealed M1 macrophages surrounding the lesion, and luminal epithelial cells lining it (Figure 6e), highlighting a preserved role of M1 macrophages in eutopic and ectopic endometrium (Figures 6c,d). Applying scCellFie to our Visium data, we detected increased metabolic activity where the luminal epithelial cells were predicted to reside. Notably, mevalonate synthesis–a metabolic marker of these cell types (Figure 4c)–scored highly in that region. Similarly, other dysregulated tasks identified with the single-cell data in luminal epithelial cells (Figure 6d), such as the HMP shunt and the phenylalanine conversion, were also highly active in the area of the endometriotic lesions (Figure 6f, Supplementary Figure 6). All of this suggests a similar metabolic shift in both eutopic and ectopic endometrium in endometriosis.

### scCellFie identifies spatial patterns of metabolism in endometrial carcinoma

We next applied scCellFie to a previously published spatial transcriptomics dataset of endometrioid endometrial cancer (EEC) covering malignant and non-malignant tissue regions^93^ (tumor microenvironments) (Figures 7a,b). Leveraging spatial information of malignant/non-malignant cells (key for capturing the architecture of the tumor microenvironment^94^), scCellFie provides deeper insight, from a metabolic perspective, into tumor organization and dynamics.

**Figure 7.**
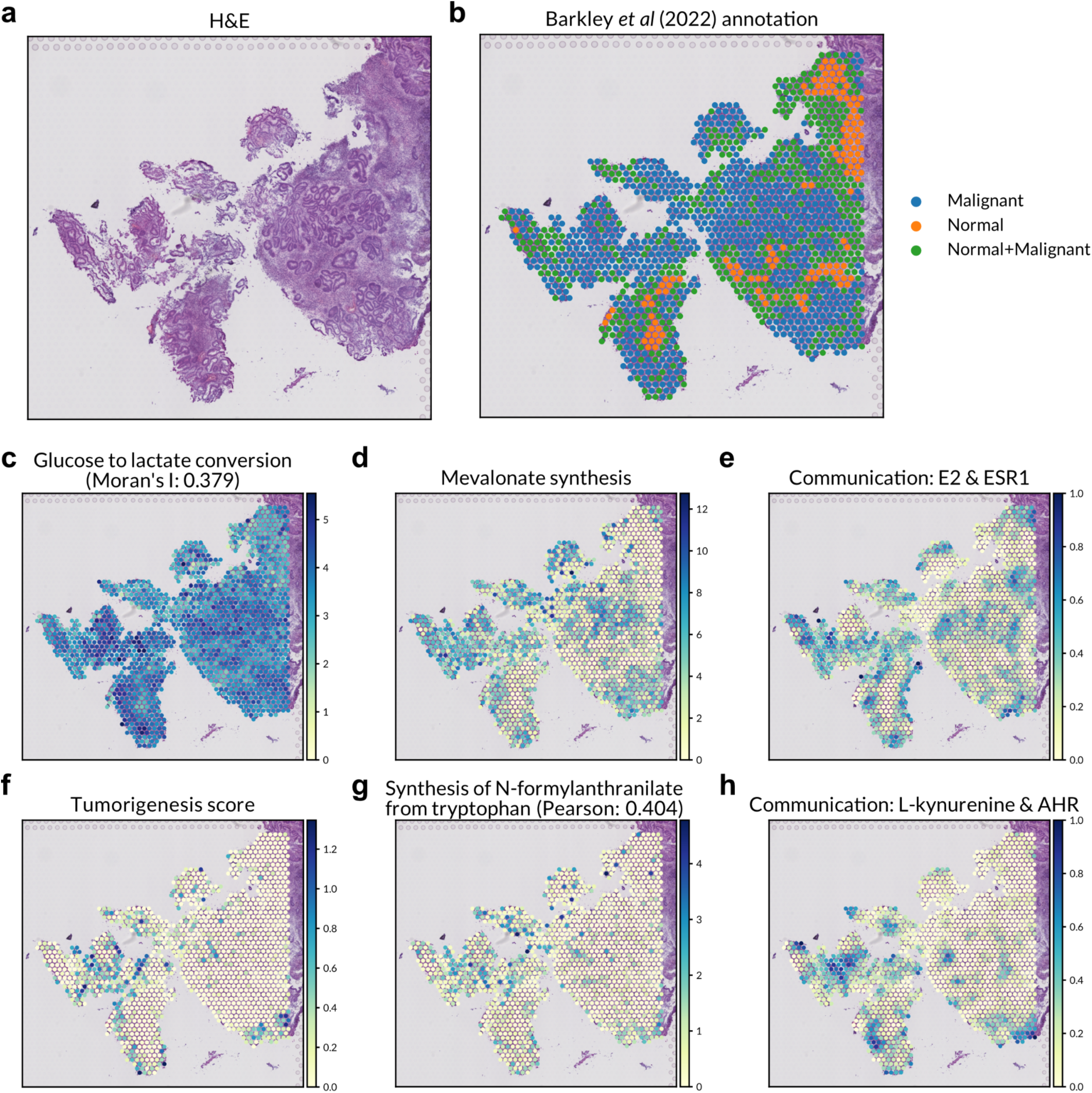
Metabolic tasks and signaling associated with endometrial tumor microenvironments. (**a**) Hematoxylin and eosin (H&E) staining and (**b**) spatial transcriptomics spots annotated for the presence of non-malignant and malignant cell types from a Visium dataset of endometrial adenocarcinoma (endometrioid endometrial cancer with squamous differentiation) from Barkley et al^93^. (**c**) Spatial organization of glucose to lactate conversion demonstrated by its Moran’s I coefficient, with a complete list of spatially organized tasks in Supplementary Figure 7. (**d**) Mevalonate synthesis identified as a metabolic marker in spots with malignant cells using scCellFie’s TF-IDF implementation. (**e**) Communication scores evaluating co-localization between estradiol (E2) biosynthesis from androstenedione and ESR1 receptor within spatial neighborhoods (0-1 scale, 1 indicating perfect co-localization). (**f**) Tumorigenesis scores were computed using Scanpy’s score genes function based on signature genes of tumorigenesis of EEC^102^ (S100A9, S100A8, LCN2, CTS1, LTF, CXCL1, SAA1, SAA2). Negative values, representing low expression across these genes, are shown as 0. (**g**) Synthesis of N-formylanthranilate from tryptophan showed the highest Pearson correlation with cancer score (in parenthesis). (**h**) CCC using L-kynurenine, an intermediate in N-formylanthranilate synthesis, and its receptor AHR within the spatial neighborhood of each spot. CCC scores were computed using the same strategy as in (**e**), in both cases using the same radius to define a neighborhood (Supplementary Figure 10b). Color bars represent corresponding metabolic, tumorigenesis, and CCC scores. EEC, endometrioid endometrial cancer; CCC, cell-cell communication.

To study if metabolic activities followed spatial patterns, we calculated the Moran’s I coefficient on scCellFie metabolic scores across all spots (**Methods**). Interestingly, glucose-to-lactate conversion (I = 0.379, Figure 7c) was one of the spatially organized metabolic tasks in endometrial carcinoma (Supplementary Figure 7), showing higher activity in regions with malignant cells (Supplementary Figure 8a). As expected, lactate production is a fundamental feature of cancer metabolism^95^. As we observed in healthy endometrium (Figure 5), non-malignant surrounding tissue also showed a basal level of lactate production (Supplementary Figure 8a).

scCellFie can help to identify region-specific metabolic activities. In contrast to the Moran’s I coefficient that is computed in a label-free manner, we can also rely on scCellFie’s marker detection strategy to find metabolic activities enriched in specific annotated regions. In this way, scCellFie identified a significant association between MVA synthesis and regions with malignant cells (Figure 7d and Supplementary Figure 8b). While our earlier analysis showed that MVA synthesis is a marker of epithelial cells in the “early secretory” phase of the healthy endometrium (Figure 4c), we now extend its relevance to malignant cells in endometrial carcinoma. This finding aligns with reported MVA pathway activation in multiple cancer types^96^ to support cell survival and growth^63^, and with the effectiveness of MVA pathway inhibitors showing antitumor activity in other cancers like ovarian carcinoma^97^.

Dysregulated estrogen synthesis has long been suggested to contribute to endometrial carcinoma as obesity is a critical risk factor of this cancer, likely by increasing estrogen production by adipose tissue^19^. However, some studies have proposed that endometrial tumors can locally produce estrogen^98,99^. To investigate this, we evaluated if regions with malignant cells can propagate the predicted increase in the synthesis of MVA–a precursor for cholesterol synthesis, which in turn serves as the main precursor for steroid hormones (Supplementary Figure 3a)–to convert cholesterol to sex hormones. We did not observe this activity (Supplementary Figure 9a), but instead scCellFie inferred activity in the intermediate step of estrogen synthesis from androgens (Supplementary Figures 9b-d). Given that androgens can be uptaken from circulation and locally converted into estrogen^100,101^, our results suggest that this may be the case in endometrial carcinoma. To understand the significance of this potential local production, we further analyzed estrogen-mediated CCC. scCellFie revealed increased co-localization of estradiol production with its receptor ESR1 in malignant regions (Figure 7e, Supplementary Figure 8c). Thus, our findings support local androgen-to-estrogen conversion within the tumor microenvironment^101^, which may contribute to this excess of estrogen signalling and, thereby, to cancer development.

To assess whether scCellFie’s inferred metabolic activities can inform risks of cancer development, we calculated the correlation between scCellFie predictions and an EEC-associated tumorigenesis score. This score summarizes the expression of 7 signature genes involved in early tumorigenesis of EEC^102^ within each spatial spot (Figure 7f). As expected, higher tumorigenesis scores were predicted for areas containing malignant cells (Supplementary Figures 8d). Among metabolic tasks, the synthesis of N-formylanthranilate from tryptophan strongly correlated with tumorigenesis potential (Figure 7g), being significantly elevated in regions containing malignant cells (Supplementary Figure 8e) due to its overlap of high activity with tumorigenesis-associated regions. This finding is particularly notable as tryptophan conversion to N-formylanthranilate is part of the kynurenine pathway^68^, which is consistent with high levels of kynurenine previously detected in endometrial cancer patients^103^. scCellFie spatial CCC analysis further revealed significantly increased co-localization between L-kynurenine synthesis and the expression of its receptor AHR in malignant regions (Figure 7h and Supplementary Figure 8f). Altogether, these results reveal a strong association between tryptophan conversion to N-formylanthranilate, the kynurenine-AHR signaling axis, and regions with high tumorigenesis potential, suggesting this metabolic task may be an important feature of endometrial carcinoma development.

## Discussion

Here we present scCellFie, a computational framework that estimates metabolic activities from transcriptomic data at true single-cell and spatial resolution. Our tool stands out from other methods by being capable to analyze large single-cell atlases, such as the CELLxGENE human cell atlas data, capturing functional relationships of enzymes through GPR rules, utilizing metabolic tasks for interpretable results, and including analysis modules to identify metabolic markers, condition-specific changes, cell-cell communication, and spatiotemporal patterns (Figure 1b). We applied it to the endometrium and revealed key metabolic activities that support both physiological functions and disease states. These include mechanisms that counteract inflammation and oxidative stress, and insights into how these processes are disrupted in endometrial disorders and the ectopic lesion microenvironment. Overall, scCellFie is a powerful and comprehensive method for identifying cellular communities driving specific metabolic functions.

scCellFie advances single-cell metabolic inference by offering a customizable and versatile framework suited to diverse research needs. Users can tailor analysis to their specific interests, either subsetting scCellFie’s predefined tasks or adding new tasks encompassing additional hormones or metabolites not present in scCellFie’s database. Furthermore, while we provide precomputed expression thresholds across the CELLxGENE atlas (Supplementary Figure 2), researchers can calculate technology-or study-specific thresholds to better reflect their experimental setups. To address data sparsity, scCellFie includes an optional KNN-based algorithm to smooth gene expression and supports interoperability with external tools such as MAGIC for missing data imputation^104^ and metacell-based methods for gene expression aggregation^105,106^. scCellFie also enables cell-cell communication analysis via curated metabolite–receptor interaction databases^107–109^, linking the potential of metabolite biosynthesis in sender cells to receptor expression in receiver cells. These features make scCellFie a flexible platform that overcomes key limitations of existing methods while providing robust, biologically meaningful interpretation of cellular metabolism.

We demonstrated the power and scalability of scCellFie’s by analyzing large-scale datasets encompassing millions of cells, resulting in a publicly available resource that maps metabolic activities across diverse human cell types (https://www.sccellfie.org). This atlas includes 2,195 cell type–organ combinations, greatly expanding upon previous human metabolic atlases, which are limited to bulk tissue resolution^48^ or a small number of cell types^110^. We envision that this resource will be a game changer by helping contextualize new data within the broader landscape of cellular metabolism across organs. For example, researchers can use our platform to determine whether a metabolic activity in their dataset is relatively high or low compared to the reference atlas, or to identify other biological aspects that are often neglected in single-cell atlases.

Beyond this global metabolic atlas, we used scCellFie to interrogate the metabolic landscape of the human endometrium, a highly regenerative tissue where tightly regulated metabolism is essential, and metabolic dysfunction is closely linked to disease development. Fine control of metabolic activity was predicted in endometrial epithelial cells, with MVA and kynurenine pathways potentially supporting proliferation and mitigating oxidative stress, respectively. Additionally, ATP generation via lactate production and estrogen bioavailability modulation may contribute to epithelial proliferation. In contrast, the metabolism of endometrial stromal cells appears to sustain tissue integrity and support uterine receptivity through metabolic tasks such as phenylalanine conversion and calnexin/calreticulin cycle, which are involved in decidualization.

We uncovered disease-associated metabolic reprogramming in the endometrium by integrating public single-cell and spatial transcriptomics datasets of endometriosis and endometrial cancer alongside newly generated spatial data. Our analysis revealed dysregulation in carbohydrate, lipid, and amino acid metabolisms, such as the production of lactate, fatty acids, and glutamine in endometriosis, which may drive atypical proliferation and proinflammatory phenotypes of epithelial cells and macrophages in this disease. This metabolic characterization highlights potential opportunities for designing or repurposing drugs that target these pathways to mitigate disease-related metabolic changes^87^. In endometrial carcinoma, malignant cells showed distinct metabolic signatures, including increased glucose-to-lactate conversion, dysregulated kynurenine pathway, and enriched intercellular communication though kynurenine-AHR and estrogen-ESR1 signaling. Notably, metabolic profiles revealed with transcriptomics data showed a distinctive spatial organization of metabolic activities within the tissue, highlighting the value of including a spatial component in the tool’s capabilities. Together, these insights underscore scCellFie’s potential to uncover new therapeutic targets by capturing the complex metabolic underpinnings of endometrial diseases.

Limitations of scCellFie include its dependency on the metabolic tasks defined in the database, which may miss other important functions or incorrectly predict activities due to misannotations. Similar to other transcriptomics-based tools, it assumes that gene expression levels closely reflect metabolic activity, which may not always occur due to differences between mRNA and protein abundances^111^, or post-transcriptional regulation and enzyme kinetics^112^. Nevertheless, scCellFie can quickly help understand metabolism under different conditions, while its predictions can be used as hypothesis generators or further refined by integrating information from different omics technologies. As such, scCellFie can readily use proteomics data, as long as proper thresholds are provided (Figure 1c), to infer metabolic activity with better confidence as enzyme presence is guaranteed with this kind of data.

Finally, scCellFie’s customizable and modular design encourages community-driven improvements, allowing researchers to contribute new metabolic knowledge (e.g. technology-specific thresholds and additional metabolic tasks) and expand its analytical capabilities with additional modules (e.g. network visualization of metabolic tasks and CCC). As a result, scCellFie stands as a dynamic platform for advancing our understanding of metabolism in health and disease.

## Methods

### Transcriptomic data and processing

scRNA-seq data was obtained from previous studies for ovaries^50^, in vivo samples of endometrium with and without endometriosis^60^, and in vitro endometrial organoids^59^. Additionally, both single-cell and single-nucleus data were obtained from the CZI CELLxGENE human and mouse cell atlases^27^. Similarly, Visium data was obtained from previous studies for human endometrium during mid secretory^59^ (A30, 152811 slide) and endometrial carcinoma^93^. Visium data of endometriotic lesions were generated with this work, and are available upon reasonable request.

Raw matrices of scRNA-seq and Visium data were library-size normalized (i.e. divided by the total number of detected RNAs in each cell/spot) and multiplied by a scale factor of 10,000, resulting in expression values of UMI counts per 10,000 (CP10K).

To avoid mis-annotated cells in the CZI CELLxGENE human cell atlas, we excluded cell types with less than 50 cells in a given organ.

### Gene expression smoothing

To address the inherent sparsity and technical noise of scRNA-seq data, scCellFie includes an optional k-nearest neighbor (KNN) smoothing approach^113^. Alternatively, external tools can be used for data imputation. The implemented smoothing relies on a pre-computed KNN graph, where each cell/spot is connected to its most similar counterparts based on their expression profiles. Then, a smoothing matrix *S* is built from the adjacency matrix summarizing the KNN graph, using weighted connections (1/k for neighbors, 0 otherwise; where k is the number of neighbors used to build the graph). Lastly, the smoothed gene expression (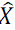) is computed as a weighted combination of a cell/spot’s original expression (left component) and the average expression of its neighbors (right component):

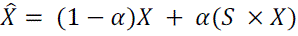

Where 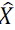 represents the smoothed expression matrix, *X* is the original expression matrix, *S* is the smoothing matrix, α controls the degree of smoothing, and × represents a matrix multiplication operation. The α parameter helps to give more importance to each cell/spot own expression levels or to the average expression of their neighborhood. A value of 0 means that no smoothing is performed, while a value of 1 transforms each cell/spot completely on their neighbors.

To manage memory constraints with large datasets, the gene expression smoothing can be performed in chunks, processing subsets of cells sequentially. These chunks can contain only cells/spots belonging to a specific cell type or cluster, or a fixed number of randomly selected cells. For each chunk, we identify all neighbors of the cells in the chunk, and compute the smoothed expression values using the chunk’s KNN graph.

Before running scCellFie, all scRNA-seq datasets analyzed in this work were smoothed using the KNN-based approach, with a number of k=10 neighbors, α parameter set to 0.33, and by running the chunk-based strategy, where each chunk represented a different cell type. In contrast, Visium datasets were smoothed by obtaining the KNN graph from the entire dataset simultaneously.

### Threshold calculation

A key input of scCellFie is a set of threshold values used to compute metabolic gene scores (Figure 1c, step 1). To assign a threshold to each metabolic gene in the GEMs Recon2^114^ and Human1^48^, we analyzed gene expression data from the April 2024 snapshot of the CELLxGENE’s human cell atlas. In this case, we adapted a previously defined local thresholding approach with a lower and an upper bound to discriminate metabolic genes that could be considered as “inactive” or “active”^46^, respectively (Figure 1c).

We first calculated the non-zero mean expression for each detected metabolic gene using CP10k-normalized values. Here, we included only cells not associated with a disease (i.e. those with “disease” metadata column labeled as “normal”), totaling ∼30 million cells. Importantly, we excluded zeros due to data sparsity, which could greatly affect how scCellFie considers a metabolic gene as “active” or “inactive” (Figure 1c). Additionally, this helped us to identify gene activities that were specific to a few cells, and that otherwise would have been masked by an excess of zeros.

Based on a previous benchmark assessing thresholding decisions^46^, we then computed the 25th and 75th percentiles of the overall distribution of non-zero expression values across all metabolic genes (Supplementary Figure 2), and assigned them as the lower and upper bounds of the local thresholding approach, respectively. This means, if a gene’s non-zero mean expression was below the 25th percentile of this distribution, its threshold was set to the 25th percentile; if it exceeded the 75th percentile, the 75th percentile was assigned. If the non-zero mean fell within these boundaries, it was used directly as the threshold (Figure 1c, step 1).

Exactly the same procedure was performed to find threshold values for mice. In this case we relied on the GEMs iMM1415^115^ and Mouse1^49^, and used mouse data from CZI CELLxGENE, totaling ∼5 million cells (July 2024 snapshot). For human and mouse thresholds, we analyzed the expression distribution (Supplementary Figure 2) and computed the non-zero mean and percentiles for each gene in a memory-efficient manner by scanning through chunks of data using the “backed” mode to read h5ad files with Scanpy^45^.

### scCellFie’s metabolic database

The metabolic database included in scCellFie has two components: 1) Pre-computed thresholds using the CELLxGENE cell atlases for humans and mice, separately, and 2) metabolic tasks, including their associated genes, reactions, GPR rules, and major category annotations. This database, for human and mouse, is available online at https://github.com/earmingol/scCellFie/tree/main/task_data.

The pre-computed thresholds were obtained as indicated in ***Threshold calculation***, while the metabolic tasks were defined using prior knowledge from different sources. Specifically, seven core functions including amino acid, carbohydrate, energy, glycan, lipid, nucleotide, and vitamin and cofactor metabolism were obtained from the original database in CellFie^39^. Five extra functions related to protein secretion, including processing in the endoplasmic reticulum, processing in the Golgi, proteostasis, translocation, and vesicle trafficking were obtained from the secretory CellFie database^47^.

We further corrected some of the reaction mappings and removed task redundancies in these sources: 1) We removed the Glucosaminyl-acylphosphatidylinositol to deacylated-glycophosphatidylinositol (GPI)-anchored protein task since it was redundant with the GPI-anchor task. 2) We corrected the step converting tyrosine to L-DOPA in the tyrosine-to-adrenaline and tyrosine-to-dopamine tasks. Specifically, we replaced the TYRDOPO reaction, which uses the TYR gene, with the TYR3MO2 reaction to use the TH gene since TYR is mainly involved in melanin production^116,117^. 3) Phenylalanine to phenylacetyl-L-glutaminate was renamed to Phenylalanine to phenylacetate (via phenylacetaldehyde) as reactions completing the production of phenylacetyl-L-glutaminate were not in the original database due to a lack of GPR rules in the available genome-scale metabolic models.

In addition, we included two new tasks: tyrosine to melanin and synthesis of GABA. For the former, we re-utilized the TYRDOPO reaction, already contained in the database, and added two more reactions to this task; TYRASE and DOPACHROMASE, which are linked to the genes TYRP1 and DCT, respectively. For the synthesis of GABA, we re-utilized the reactions ABTArm and GLUDC. All reaction IDs here are present in the GEM Recon1^118^, except DOPACHROMASE that was manually added.

Finally, we added a new category of tasks to capture the synthesis of non-peptide hormones (Figure 2b), in this case based on prior knowledge in the GEMs Human1^48^ and Mouse1^49^. Specifically, we defined seven new tasks associated with the steroid hormones testosterone, progesterone, and estrogens (Supplementary Figures 3a,b). After manually curating all the information collected from the distinct sources, the scCellFie database resulted in 218 metabolic tasks for humans and 203 for mice.

### scCellFie’s metabolic score

scCellFie infers metabolic task scores from transcriptomic data through a multi-step process that integrates gene expression in individual cells with metabolic network knowledge (Figure 1c). Starting from single-cell or spatial expression profiles, scCellFie first processes the data through gene expression normalization (CP10k values) and optional smoothing based on cell neighborhoods.

The first step of predicting metabolic activity consists of computing gene scores using a formula previously implemented in CellFie^29^:

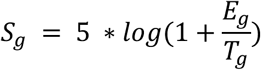

where *S_g_* is the assigned gene score for a metabolic gene *g*, and *E_g_* and *T_g_* are the expression level and threshold for that gene, respectively. Log here represents the natural logarithm.

The second step consists of linking gene scores in a functional way through GPR rules that define the relationships between genes, enzymes, and metabolic reactions according to Boolean logic (AND/OR operations). For each reaction and its associated GPR rule, the minimum gene score is selected for AND operations, and the maximum score for OR operations, reflecting the biological requirement that all protein subunits are necessary in protein complexes (AND operation) while isoenzymes can substitute each other (OR operation). In this case, scCellFie handles AND/OR operations on gene scores through the COBRApy^119^ package. The resulting score after AND/OR operations from the GPR rule corresponds to the reaction activity level (RAL). RAL normally corresponds to the gene score of the reaction’s determinant gene (i.e. the gene that dominated the AND/OR operations within the GPR rule), which is also weighted by a specificity correction factor (C) that accounts for this determinant gene participating in multiple reactions. This correction divides a gene’s score by the number of reactions it influences (i.e. the gene is determinant in those reactions), thereby preventing highly promiscuous genes from dominating the scoring. Mathematically this could be represented as:

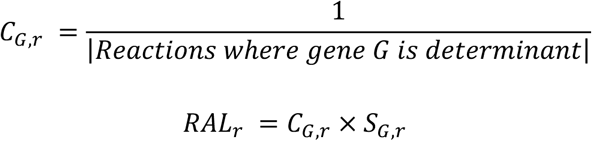

where *C_G,r_* is the specificity correction factor for the determinant gene *G* in reaction *r*, meaning that gene *G* had the dominant gene score in reaction *r* (*S_G,r_*) after performing the binary operations in the GPR for that reaction.

Finally, the third step corresponds to calculating metabolic task scores by averaging the reaction activity levels across all reactions involved in each task. The mathematical formulation is:

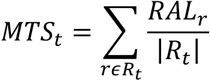

where *MTS_t_* is the metabolic score for task *t*, *RAL_r_* is the reaction activity level for reaction r, *R_t_* is the set of reactions involved in task t, and |*R_t_*| is the number of reactions in that set. Of note, MTS should not be compared across different tasks, as each task involves distinct genes and reactions. However, comparisons for a given metabolic task can be made across different cells and cell types.

### Markers selection methods in scCellFie

To identify metabolic markers, we applied the Term Frequency-Inverse Document Frequency (TF-IDF) approach^120^ included in scCellFie, which is often employed in the natural language processing field to identify important words (metabolic tasks) in a document (cell type). This strategy was implemented as in the SoupX package^121^. Furthermore, considering scCellFie integration with the Scanpy ecosystem^45^, we also applied a logistic regression approach implemented in Scanpy^122^. Additional filters to select relevant markers where applied based on each method’s score distribution.

### Statistical tests and differential analysis

To compare metabolic scores of a given task across different cell types, annotated spatial spots, or disease conditions, we performed a Wilcoxon rank-sum test. To account for multiple comparisons, P values were adjusted using the Benjamini-Hochberg procedure, controlling the false discovery rate (FDR). Metabolic tasks were considered significant if FDR < 1%. In addition to statistical significance, we assessed the magnitude of differences using Cohen’s D, a standardized effect size measure. To ensure biologically meaningful differences, we applied a selection criterion of |Cohen’s D| ≥ 0.75.

To reduce sample size bias (or achieve more statistical power) when performing the differential analysis for endometriosis, we only tested cell types that have at least 170 single cells per condition, and cell types that did not have a number of cells more than 64 times greater in one condition than the other.

### Generalized additive models

To identify metabolic activity patterns across predefined cell-type trajectories, we applied Generalized Additive Models (GAMs) to the metabolic task scores predicted by scCellFie. For each metabolic task, we fitted a GAM with the following formula:

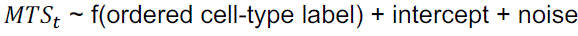

where *MTS_t_*, representing the metabolic score of task *t*, is the response variable and f(ordered cell-type label) is a smooth function of the cell-type trajectory. The cell types were encoded numerically based on their position in our predefined ordered list. The model was implemented using a normal distribution with identity link function. We represented the smooth function using penalized splines with 10 basis functions (n_splines=10) of order 3 and a smoothing penalty strength of 0.6 (lam=0.6), providing sufficient flexibility to capture non-linear patterns while preventing overfitting. We applied the GAMs directly to individual cell data without pseudo-bulk aggregation, maintaining single-cell resolution in our analysis.

To select metabolic tasks with clear trends across cell-type trajectories, we filtered the results using two criteria: McFadden’s adjusted R^2^ > 0.2 and scale parameter > 0.2. In this way, we prioritized metabolic tasks whose GAMs predict better than a null model (McFaddenn’s adjusted R^2^) and with sufficient variation across cell type labels to be biologically meaningful (scale parameter). GAMs were implemented as part of the scCellFie tool using the pyGAM package^123^.

### Identification of spatially organized metabolic tasks

To analyze the spatial organization of metabolic tasks within tissues, we employed Moran’s I spatial autocorrelation statistic, as implemented in Squidpy^124^. After computing cell-specific metabolic task scores using scCellFie, we calculated Moran’s I for each metabolic task across the spatial coordinates of Visium spots. This statistic quantifies whether metabolically similar cells tend to cluster together (positive Moran’s I values) or disperse in alternating patterns (negative values), with values near zero indicating random spatial distribution.

### Cell-cell communication analysis

Metabolic tasks involved in the synthesis of specific metabolites can contribute to CCC. To quantify these interactions, distinct CCC scores can be computed by integrating metabolic activity in sender cells with receptor expression in receiver cells^125,126^. By combining scCellFie-derived metabolic scores, receptor gene expression, and prior knowledge of metabolite-based ligand-receptor interactions^107^ (e.g. kynurenine-AHR and estradiol-ESR1), we computed CCC scores for both single-cell and spatial transcriptomic data.

For single-cell data, we first aggregated metabolic scores and receptor gene expression at the cell type level by averaging values across individual cells. We then computed the CCC score as the geometric mean of the metabolic score for metabolite synthesis in the sender cell type and the expression of the cognate receptor in the receiver cell type. To identify biologically relevant interactions, we applied the following filters: (1) over 10% of cells in the sender population must have a metabolic score above natural log(2), (2) over 10% of cells in the receiver population must express the cognate receptor, and (3) the CCC score must be at least one standard deviation above the mean CCC score for the same ligand-receptor pair across all cell type pairs.

For spatial data, we defined a co-localization score per spot, based on each spot’s interactions with its neighbors, referred to as ‘pairwise_concordance’ in the tool. This score was computed by first identifying neighboring spots within a predefined radius (Supplementary Figure 10). Each sender-receiver spot pair within the neighborhood, including self-interactions, was then evaluated using a binary strategy^125^, wherein metabolic activity and receptor expression had to surpass defined thresholds. These thresholds were set as the mean metabolic score and receptor expression across all spots, respectively. Finally, the co-localization score was determined as the fraction of spot pairs meeting these criteria within a given neighborhood.

In both cases, gene expression of the receptors was used as log1p(CP10k).

### Location of cell types in Visium data

To localize cell types in the Visium transcriptomics slides, we used the cell2location tool^92^ with default parameters. As reference, we used the HECA single-cell dataset with its original annotations. Thus, we could identify epithelial and immune cells in the slides of ectopic lesions. Corresponding plots represent estimated abundance of cell types (cell densities) in Visium spots.

## Data availability

scRNA-seq data for ovary is publicly available in GEO under accession number GSE192722. scRNA-seq data for the HECA (in vivo data) and endometrial organoids (in vitro data) are publicly available in ArrayExpress under accession numbers E-MTAB-14039 and E-MTAB-10283, respectively. Processed and annotated matrices can be accessed and downloaded from www.reproductivecellatlas.org. Visium data for the secretory endometrium is publicly available in ArrayExpress under accession number E-MTAB-9260. Visium data for endometrial carcinoma is publicly available in GEO under accession number GSE203612. Visium data of endometriotic lesions were generated with this work, and are available upon reasonable request.

## Code Availability

scCellFie and its database are publicly available in the tool’s GitHub repository (https://github.com/earmingol/sccellfie). Code and Jupyter Notebooks used in this work’s analyses have been deposited to a GitHub repository (https://github.com/ventolab/scCellFie-Paper-Code).

## Acknowledgements

We acknowledge the support of Cellular Genetics IT, New Pipeline Group and DNA pipelines of the Wellcome Sanger Institute. We acknowledge funding from Wellcome Leap’s Dynamic Resilience Program (jointly funded by Temasek Trust). E.A. is supported by the European Commission through a Marie Skłodowska-Curie Actions Postdoctoral Fellowship (Project 101208051). This work was also supported by funding from NIH (R35 GM119850; N.E.L.). The authors thank Aidan Maartens for insightful feedback.

## Author contributions

E.A. and R.V.T. conceived the work. L.G. and N.E.L. provided important insights about the analyses and the method. E.A. implemented scCellFie, curated the database, and performed computational analyses across all datasets. J.A. curated the database and analyzed scRNAseq data of ovaries. M.P. downloaded and processed the CZI CELLxGENE human cell atlas. E.A. and M.P. implemented the online website for visualizing activity of metabolic tasks from CZI’s atlas or each user’s results. M.Mareckova, C.I.M., C.B., and K.Z. processed the tissue sample and generated the Visium data of the ectopic lesion (endometriosis).

V.L. prepared and analyzed data of the ectopic lesion. M.Moullet annotated cell types in datasets of endometrial diseases. E.A., J.A., and J.T.H.L. prepared the tutorials. E.A. wrote the paper, L.G. and R.V.T. revised the manuscript, and all authors carefully reviewed, discussed and edited the paper.

