## Supplementary Figures for "Atlas-scale metabolic activities inferred from single-cell and spatial transcriptomics"

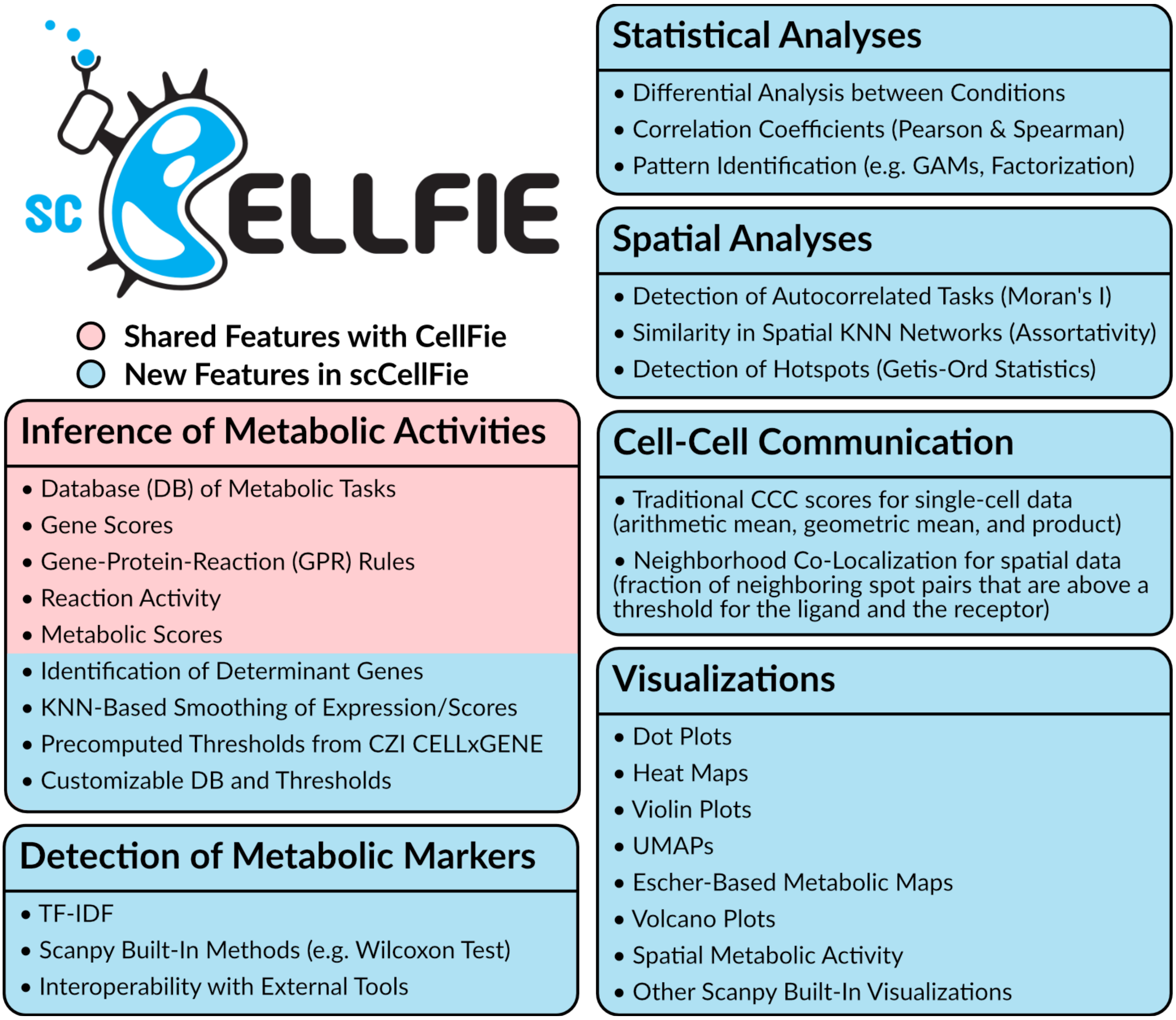


**Supplementary Figure 1. scCellFie is a comprehensive framework to study metabolism in large atlases.** By including new features such as a gene expression smoothing strategy, thresholds computed across all individual cells in the human cell atlas, and customizable database, scCellFie enables the inference of metabolic activities from single-cell and spatial data. It uses a strategy inspired by the original CellFie method (designed for bulk and small single-cell data), while incorporates multiple analysis modules that extends the tool use beyond only predictions of metabolic activity, including detection of metabolic markers, statistical tests, spatial analysis, pattern identification, and multiple visualizations. Although the original CellFie includes limited visualizations, ImmCellFie is an online tool that extends CellFie visualization capabilities, covering some of those included in scCellFie; however, it is limited to bulk data.

**a**

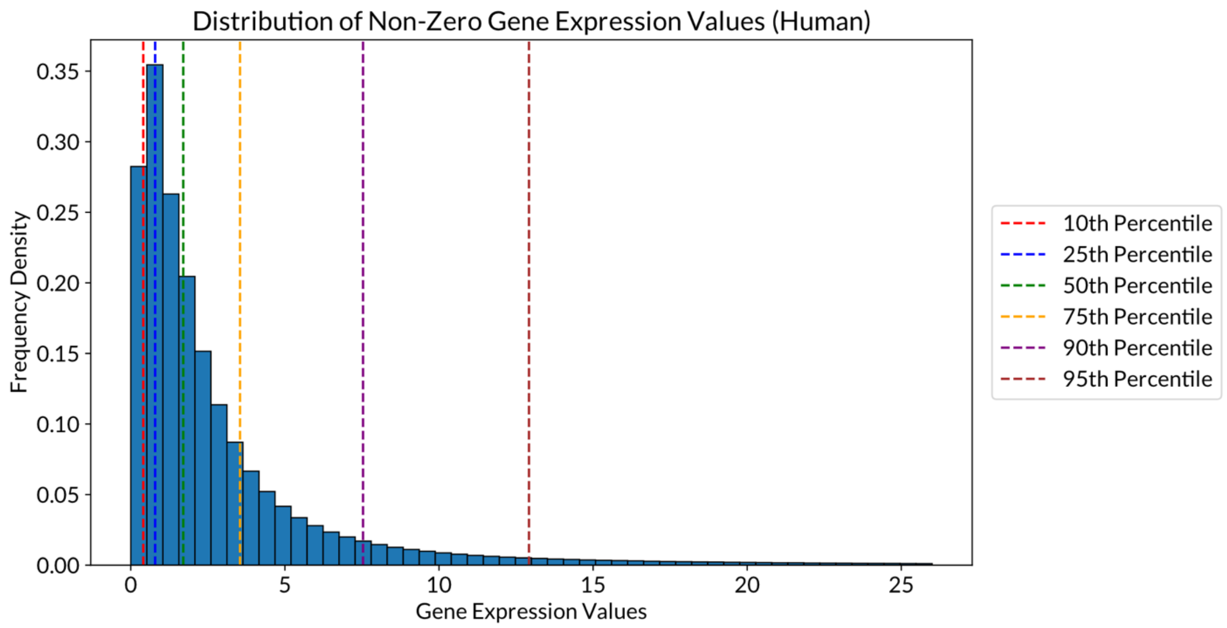


**b**

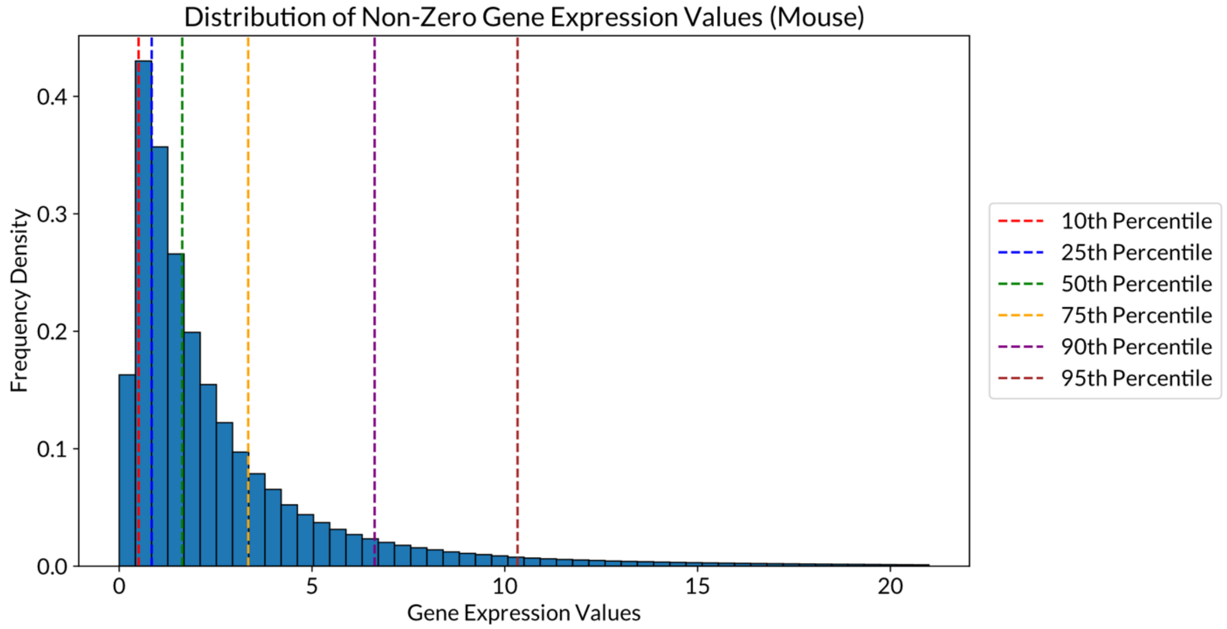


**c**

| | Lower Bound (25th percentile) | Upper Bound (75th percentile) |
|---|---|---|
| **Human** | 0.789952 | 3.544842 |
| **Mouse** | 0.842318 | 3.344482 |

**Supplementary Figure 2. Distributions of metabolic gene expression in the CZI CELLxGENE human/mouse cell atlas.** (**a**) Histogram showing the distribution of the non-zero expression (CP10k) of 3,061 human metabolic genes, obtained from the genome-scale metabolic models Recon2 and Human1, across ~30 million cells (with a "normal" label for the disease metadata) in the CELLxGENE human cell atlas (April 2024 snapshot). (**b**) Histogram showing the distribution of the non-zero expression (CP10k) of 3,025 mouse metabolic genes, obtained from the genome-scale metabolic models iMM1415 and Mouse1, across ~5 million cells (with a "normal" label for the disease metadata) in the CELLxGENE mouse cell atlas (July 2024 snapshot). Dashed lines in (**a**) and (**b**) represent the percentiles as colored in the legend. (**c**) Expression boundaries used to define the thresholds needed for scCellFie calculations. Lower and upper bounds were to the 25th and 75th percentiles in (**a**) and (**b**), respectively. Each gene threshold is calculated as the non-zero expression average across all cells, and if this mean is below the lower bound, the threshold is set to the lower bound; or if this mean is above the upper bound, the threshold is set to the upper bound.

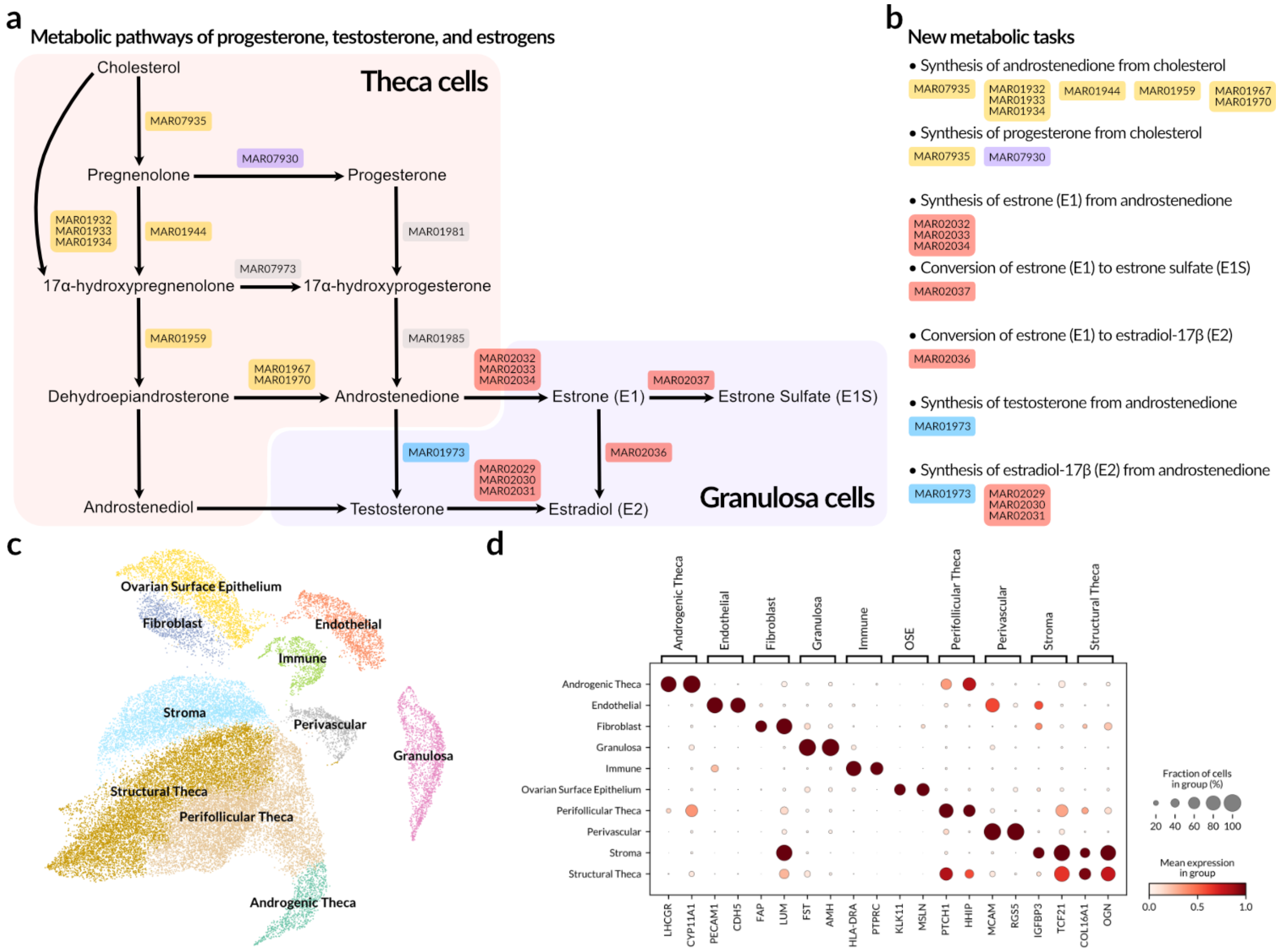


**Supplementary Figure 3. Addition of the hormones metabolism to the scCellFie's metabolic database.** (**a**) Biosynthesis network of sex hormones including progesterone, testosterone, estrone and estradiol. Colored boxes show reaction IDs (starting with "MAR") for the steps in this network that were mapped to a reaction in the genome-scale metabolic model Human1. Boxes containing multiple reaction IDs summarize multiple metabolic steps into a single step. Yellow boxes are related to the cholesterol to androstenedione conversion; purple box represents the pregnenolone to progesterone conversion, light blue box indicates the reaction transforming androstenedione to testosterone, and red boxes indicate reactions involved in the estrogen biosynthesis. Gray boxes show reactions identified in Human1 as part of the sex hormones biosynthesis but that were excluded from the scCellFie database, as they are associated with exactly the same enzymes as one of the reactions already included (e.g. MAR01981 is equivalent to MAR01944 in terms of involved enzymes). Colored shades indicate the distinct steps of these biosynthetic pathways that are performed by specialized ovarian cell types (theca and granulosa cells). (**b**) New metabolic tasks included in the scCellFie database as part of the hormones metabolism category, showing the list of reactions involved in each of them. (**c**) UMAP showing the re-annotated cells in the public scRNA-seq dataset of ovaries (Guahmich et al, 2023) used to validate the new metabolic tasks. (**d**) Marker genes distinguishing the different cell types in the dataset.

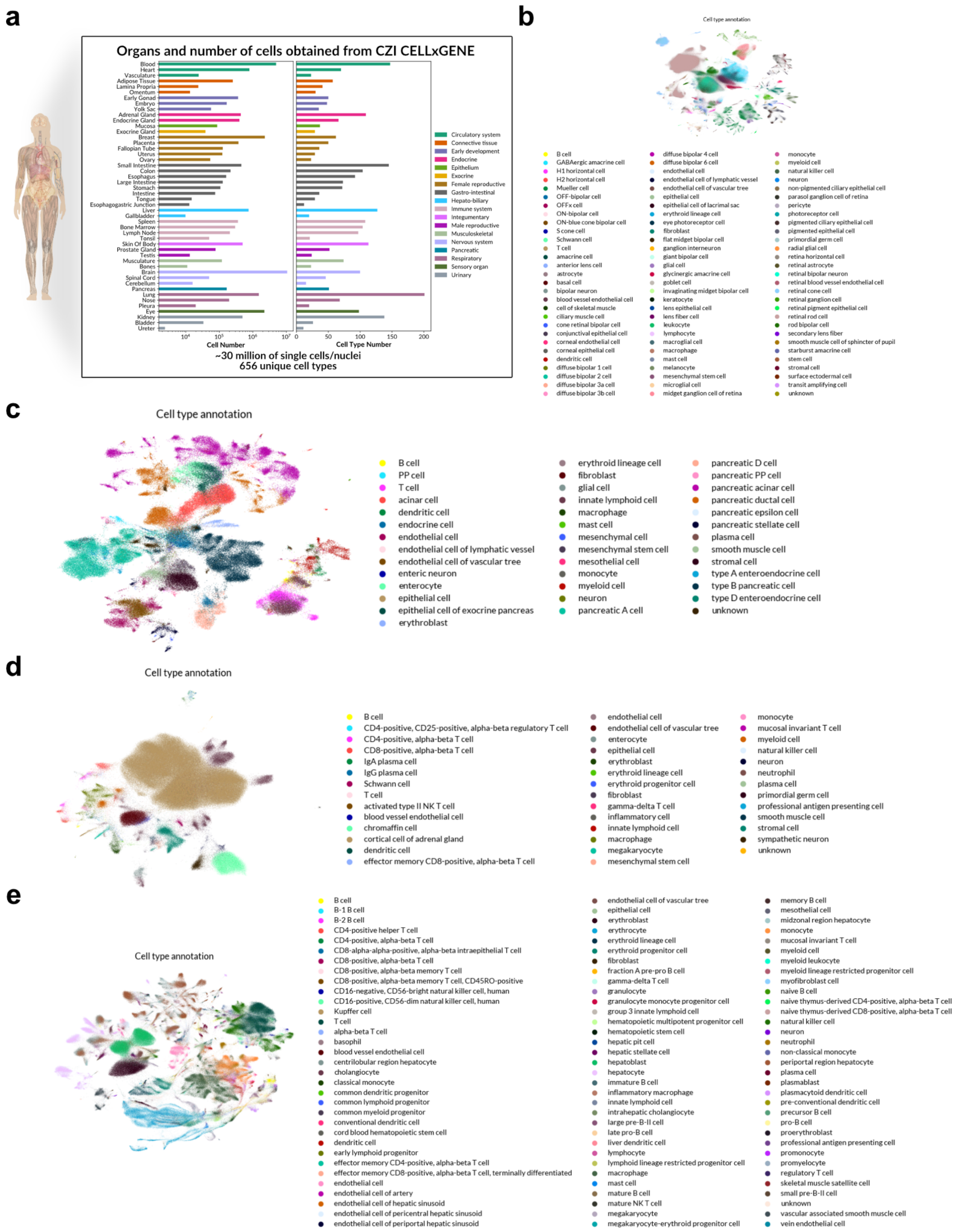


**Supplementary Figure 4. Organs and cells in the CZI CELLxGENE human cell atlas.** (**a**) Composition of the CZI CELLxGENE human cell atlas, showing the different organs and their categories, and the number of cells and cell types in each of them. (**b**-**e**) UMAPs of cells included in datasets of the (**b**) eye, (**c**) pancreas, (**d**) adrenal gland, and (**e**) liver in the CELLxGENE atlas, and their annotation based on their cell ontology. Annotations are shown as originally provided in the CELLxGENE atlas.

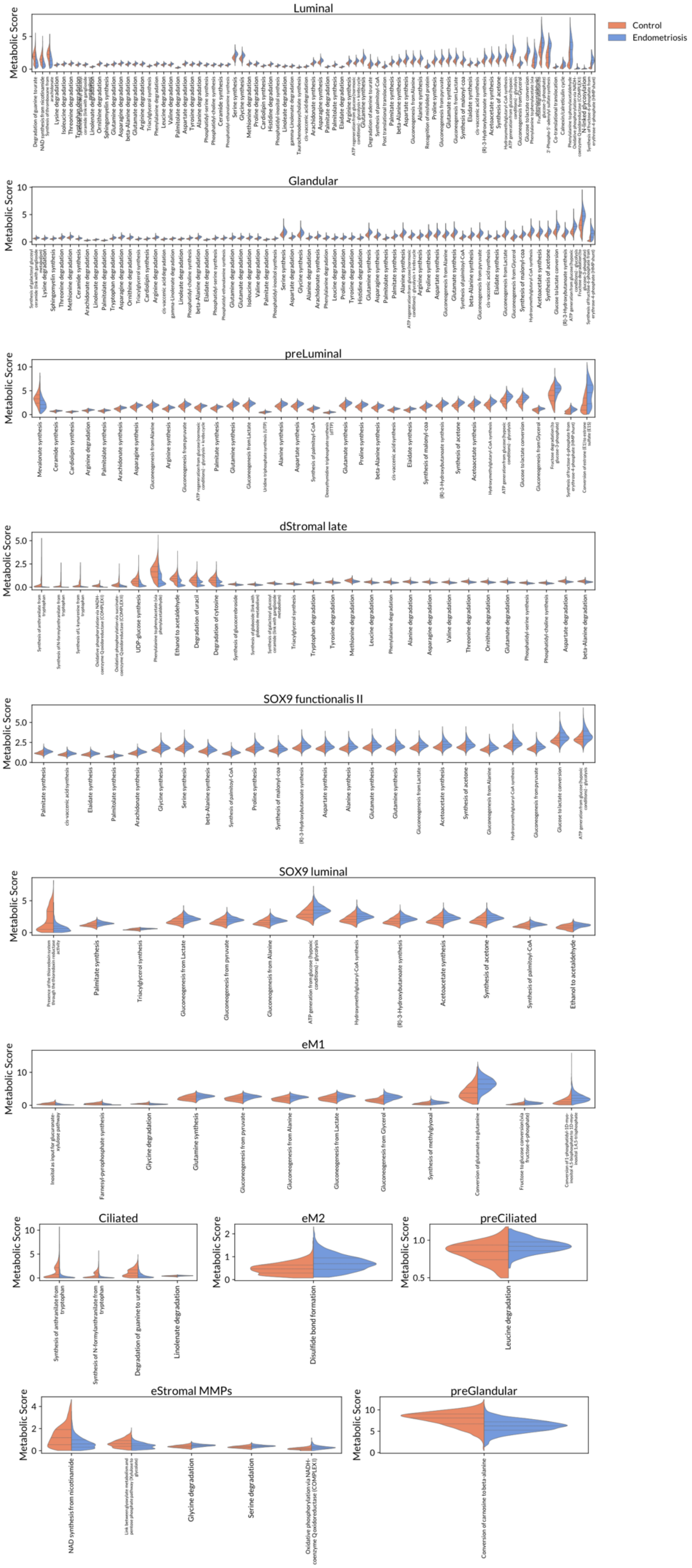


**Supplementary Figure 5. Metabolic changes in the eutopic endometrium of donors with and without endometriosis.** Violin plots are shown for cell types with at least one significantly altered metabolic task, as identified in Figure 6 of the main manuscript. Each violin plot represents the distribution of metabolic scores at the single-cell level for the corresponding cell type. Horizontal lines within each violin denote the first quartile (Q1), median, and third quartile (Q3). Only tasks with significant differences between conditions are shown (|Cohen's d| > 0.75 and FDR < 0.01, Wilcoxon rank-sum test). Colors in the legend indicate the donor groups represented in each plot.

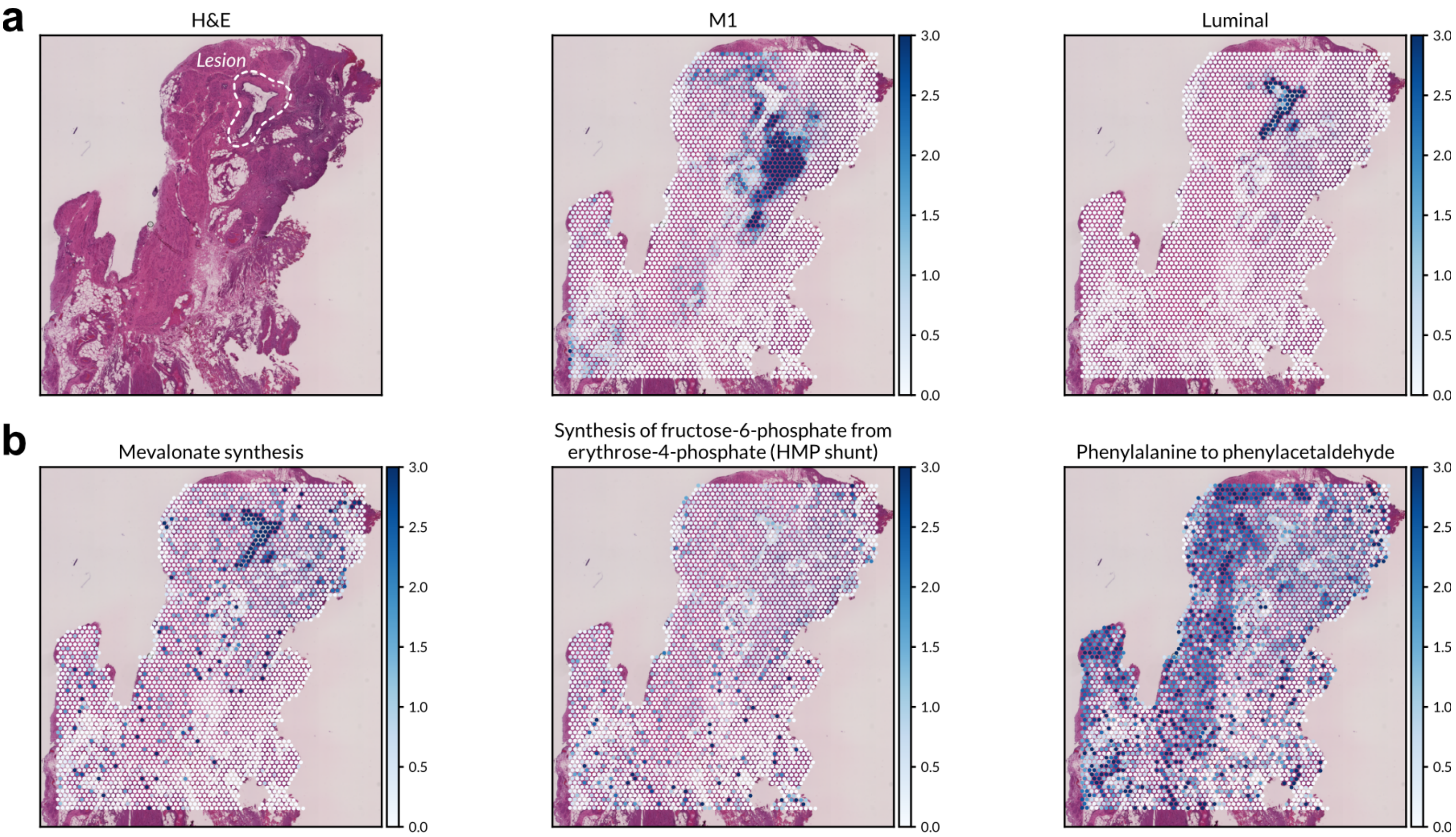


**Supplementary Figure 6. Spatial transcriptomics and metabolic activity in a superficial peritoneal lesion from a woman with endometriosis.** (**a**) Visium data from a deep infiltrating lesion of a donor with endometriosis. H&E image and cell2location scores are displayed, covering the location of M1 macrophages, and luminal epithelial cells. Dashed line encloses the endometriosis lesion in the H&E image. Cell distributions inferred with cell2location were obtained by using HECA single-cell transcriptomes as reference. Only cell types with a high score are displayed. (**b**) Metabolic scores are presented for one metabolic marker of epithelial cells (mevalonate synthesis task) and for two differentially altered tasks in luminal epithelial cells (synthesis of F6P from E4P (HMP shunt) and phenylalanine to phenylacetaldehyde). F6P, fructose-6-phosphate; E4P, erythrose-4-phosphate.

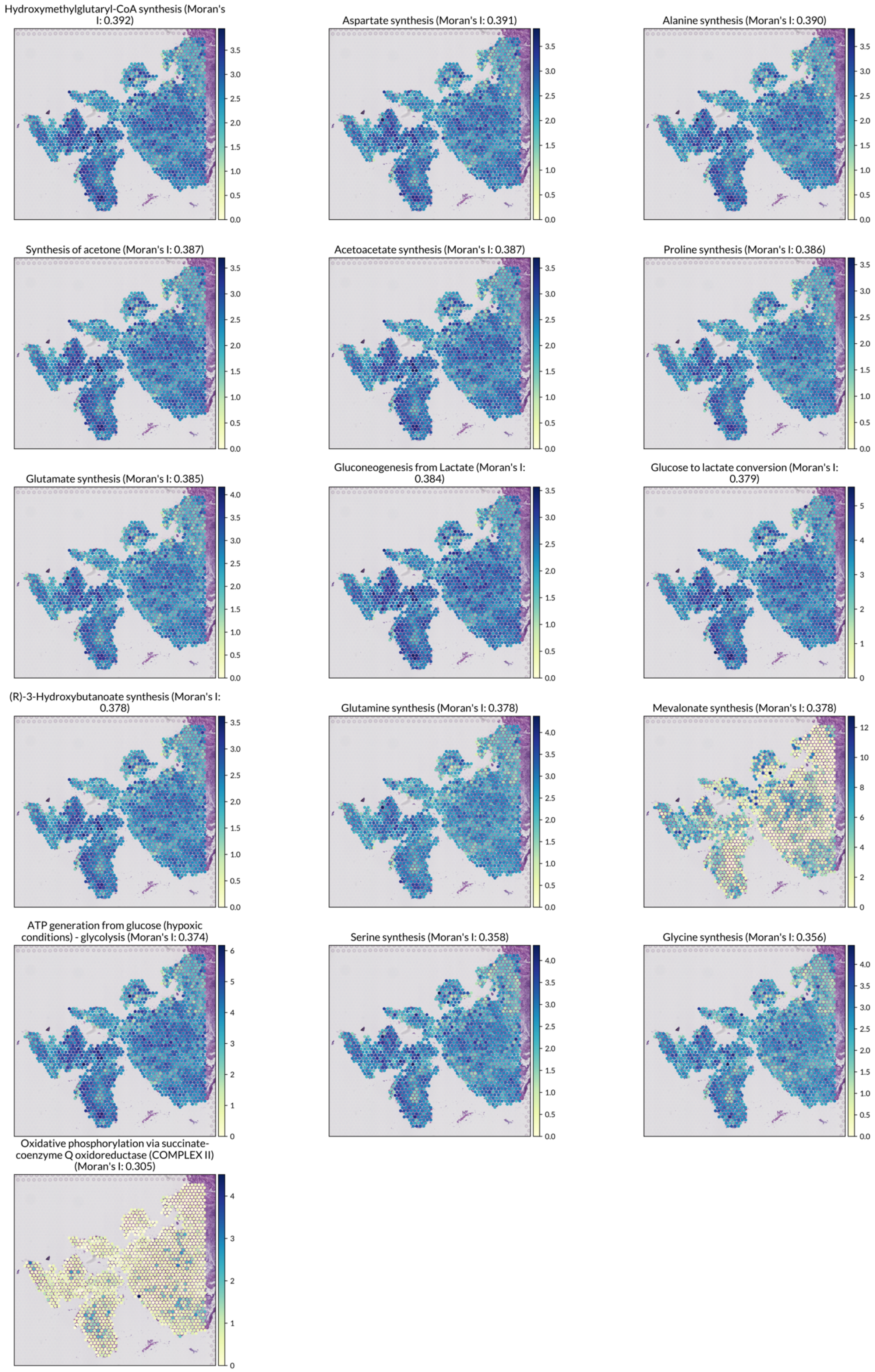


**Supplementary Figure 7. Spatially organized metabolic tasks in endometrial carcinoma.** Spatial distributions of metabolic scores are shown for tasks with a Moran's I coefficient greater than 0.3, indicating important spatial autocorrelation (i.e. spatially organized). To ensure biological relevance, tasks were further filtered to include only those with an activity score over $5 \times \log(2)$ in at least one Visium spot. This threshold indicates that all determinant genes required for the reactions comprising the task are considered "active," meaning their expression exceeds the predefined thresholds used to compute their gene scores. The Moran's I coefficient for each task is shown in parenthesis next to the task name, above its corresponding plot. This analysis was performed on a public Visium dataset of endometrial adenocarcinoma (Barkley et al, 2022).

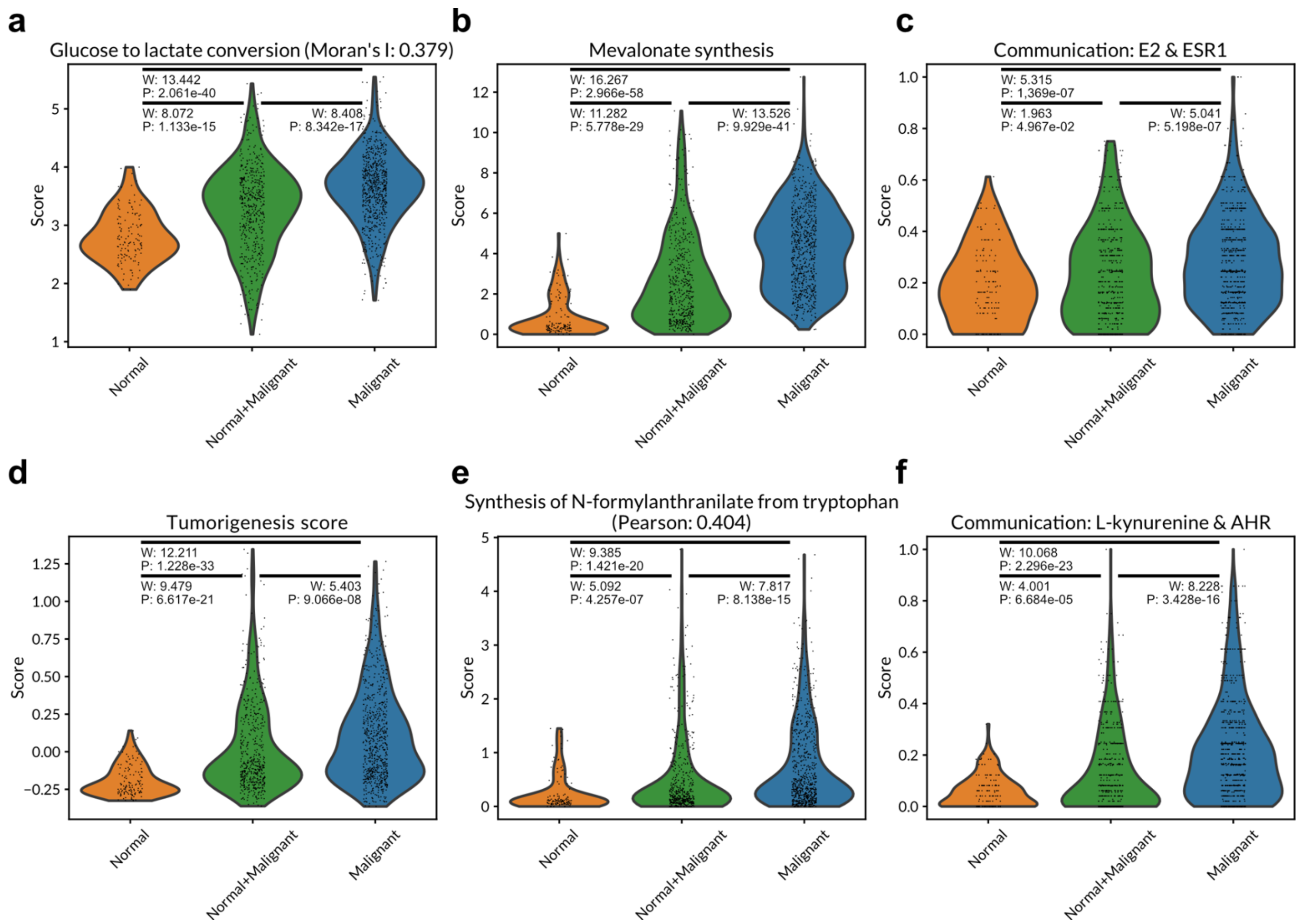


**Supplementary Figure 8. Comparison of metabolic, communication, and tumorigenesis scores in regions with and without malignant cells.** (**a**-**f**) Violin plots show the distribution of various scores, including metabolic, cell-cell communication, and tumorigenesis scores, across Visium spots, grouped according to their original labels indicating the presence of normal or malignant cells. Each panel shows the distribution of: (**a**) Metabolic scores for glucose-to-lactate conversion, a spatially organized metabolic task based on its Moran's I coefficient. (**b**) Metabolic scores for mevalonate synthesis, selected as a spatial marker distinguishing tissue regions by the presence of malignant cells. (**c**) Communication scores indicating co-localization of estradiol (E2) synthesis and expression of its receptor ESR1 across each spot's neighborhood. (**d**) Tumorigenesis scores based on the expression of genes associated with early-stages of endometrioid endometrial carcinoma (S100A9, S100A8, LCN2, CTS1, LTF, CXCL1, SAA1, SAA2). (**e**) Metabolic scores for the synthesis of N-formylanthranilate from tryptophan, the task most strongly correlated with tumorigenesis scores. (**f**) Communication scores indicating co-localization of kynurenine synthesis and expression of its receptor AHR across each spot's neighborhood, as kynurenine synthesis encompasses earlier steps in the pathway from tryptophan that ultimately lead to N-formylanthranilate.

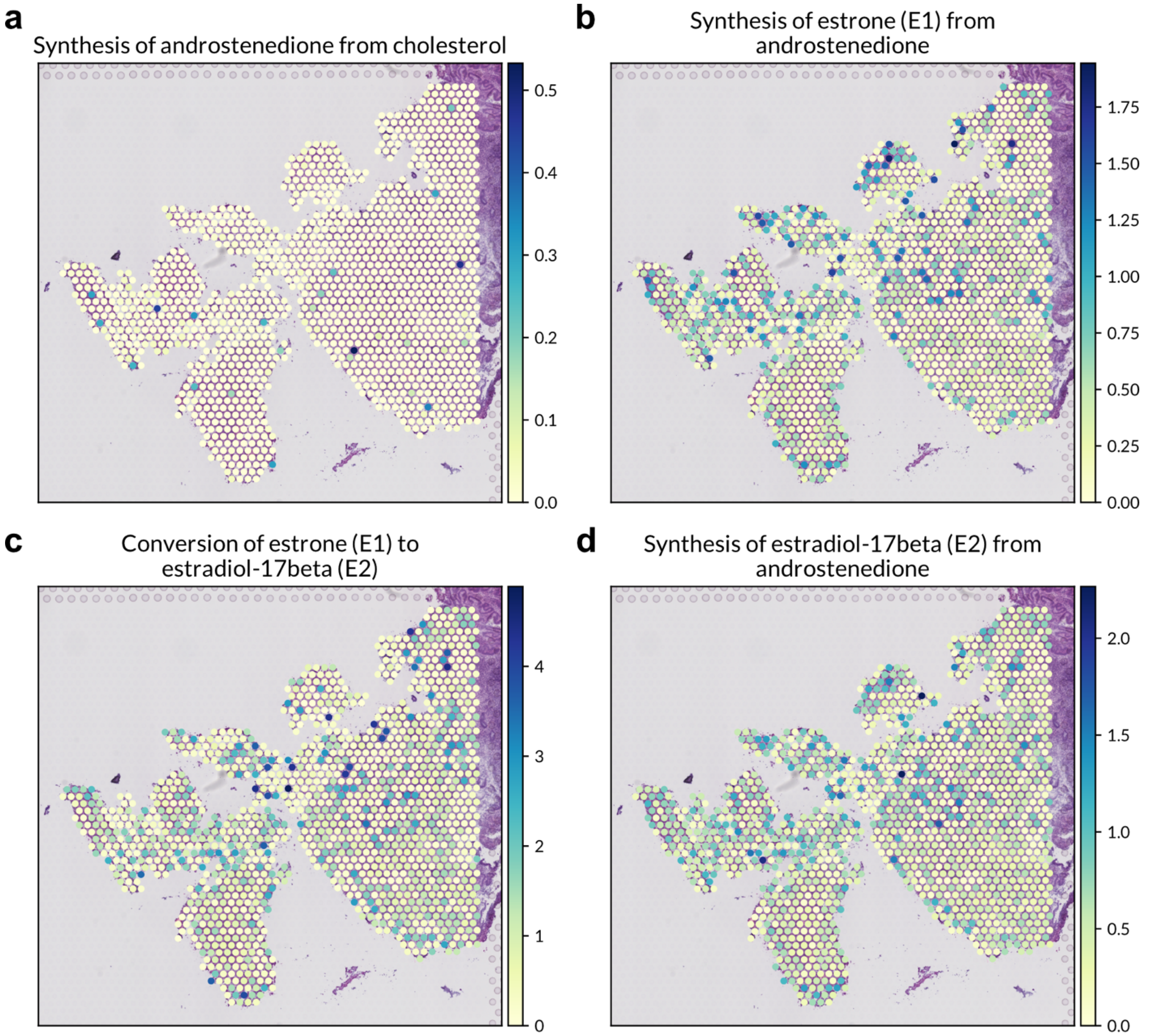


**Supplementary Figure 9. Metabolic activities associated with estrogen production in endometrial carcinoma.** Metabolic scores for tasks involved in the biosynthesis of estrone (E1) and estradiol-17β (E2) from cholesterol, inferred using scCellFie on Visium data from endometrial carcinoma (Barkley et al., 2022). The panels show the sequential steps in this pathway: (**a**) conversion of cholesterol to androstenedione, representing a rate-limiting step in estrogen synthesis; (**b**) conversion of androstenedione to E1; (**c**) interconversion of E1 to E2; and (**d**) direct conversion of androstenedione to E2.

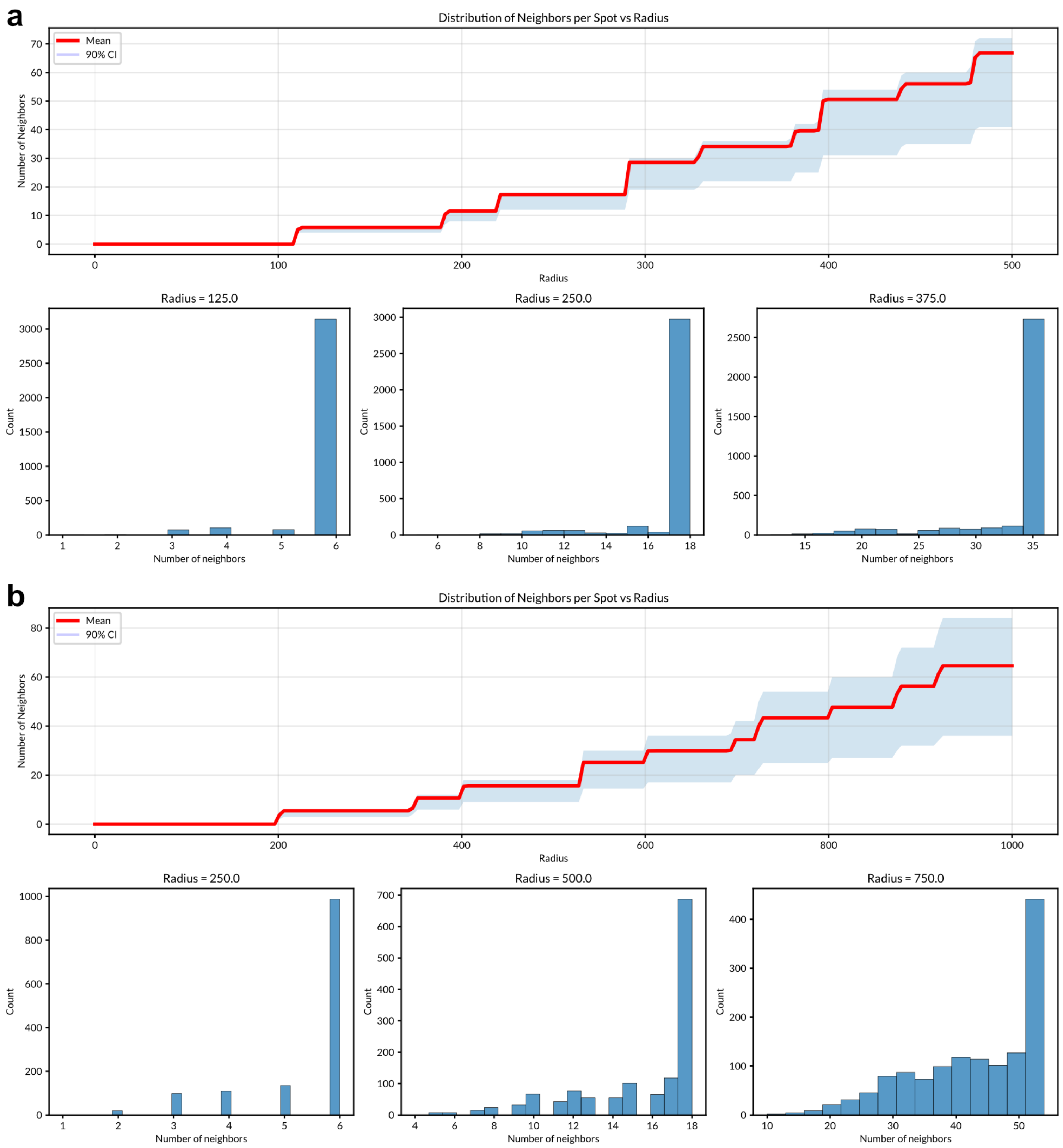


**Supplementary Figure 10. Defining spatial neighborhoods in Visium data based on spot proximity.** (**a**) Statistics for defining neighborhoods in a healthy endometrial sample. (**b**) Statistics for defining neighborhoods in an endometrial carcinoma sample. The number of neighboring spots as a function of the distance (radius) from a given spot is shown in the top panel of (**a**) and (**b**). The red line shows the mean number of neighbors across all spots for a given radius, with the light blue shade representing the 90% confidence interval. Bottom histograms in (**a**) and (**b**) display the distribution of spots according to their number of neighbors within three selected radii, where the x-axis indicates the number of neighbors and the y-axis shows the number of spots with that value. A radius of 125 and 250 were selected, respectively, for (**a**) and (**b**) to be used in downstream analyses, as it maximizes the number of spots with exactly six neighbors. In other words, this number of neighbors reflects the case where all directly adjacent (immediate) neighboring spots are included.
